# What the Zebrafish’s Eye Tells the Zebrafish’s Brain: Retinal Ganglion Cells for Prey Capture and Colour Vision

**DOI:** 10.1101/2020.01.31.927087

**Authors:** M Zhou, J Bear, PA Roberts, FK Janiak, J Semmelhack, T Yoshimatsu, T Baden

**Affiliations:** School of Life Sciences, University of Sussex, Brighton, UK; Hong Kong University of Science and Technology, HK; Institute for Ophthalmic Research, University of Tübingen, Tübingen, Germany

**Author notes:** Equal contribution (M Zhou, J Bear).

## Abstract

In vertebrate vision, the tetrachromatic larval zebrafish permits non-invasive monitoring and manipulating of neural activity across the nervous system *in vivo* during ongoing behaviour. However, despite a perhaps unparalleled understanding of links between zebrafish brain circuits and visual behaviours, comparatively little is known about what their eyes send to the brain in the first place via retinal ganglion cells (RGCs). Major gaps in knowledge include any information on spectral coding, and information on potentially critical variations in RGC properties across the retinal surface to acknowledge asymmetries in the statistics of natural visual space and behavioural demands. Here, we use *in vivo* two photon (2P) imaging during hyperspectral visual stimulation as well as photolabeling of RGCs to provide the first eye-wide functional and anatomical census of RGCs in larval zebrafish.

We find that RGCs’ functional and structural properties differ across the eye and include a notable population of UV-responsive On-sustained RGCs that are only found in the acute zone, likely to support visual prey capture of UV-bright zooplankton. Next, approximately half of RGCs display diverse forms of colour opponency - long in excess of what would be required to satisfy traditional models of colour vision. However, most information on spectral contrast was intermixed with temporal information. To consolidate this series of unexpected findings, we propose that zebrafish may use a novel “dual-achromatic” strategy segregated by a spectrally intermediate background subtraction system. Specifically, our data is consistent with a model where traditional achromatic image-forming vision is mainly driven by long-wavelength sensitive circuits, while in parallel UV-sensitive circuits serve a second achromatic system of foreground-vision that serves prey capture and, potentially, predator evasion.

## INTRODUCTION

In vertebrate vision, all information sent from the eye to the brain is carried by the axons of retinal ganglion cells (RGCs) (Masland, 2012; Wässle, 2004). Classically, RGC types are thought to encode information about image features such as the colour, speed or orientation of an edge. Through a mosaic arrangement of an RGC type across the retinal surface this information can then be transmitted for all of visual space. However, what exactly all these features are (Baden et al., 2016; Vlasits et al., 2019; Wässle, 2004), and to what extent their structure and function is truly homogeneous over the retinal surface to meet the demands of an animal’s species-specific visual ecology (Bleckert et al., 2014; Chang et al., 2013; Sabbah et al., 2017; Sinha et al., 2017; Szatko et al., 2019; Warwick et al., 2018) remains an area of active research (Baden et al., 2020). Moreover, directly linking RGC types to specific visual behaviours remains a central challenge in vision science (Baden et al., 2020; Lettvin et al., 1959).

Here, zebrafish offer a powerful way in (Bollmann, 2019). Their excellent genetic access and largely transparent larval stage has made it possible to probe their visual circuits *in vivo* while animals were performing visual behaviours such as prey capture (Antinucci et al., 2019; Bianco et al., 2011; Mearns et al., 2019; Ramdya and Engert, 2008; Semmelhack et al., 2014) or predator evasion (Dunn et al., 2016; Preuss et al., 2014; Temizer et al., 2015). In fact, prey-capture-like behaviours can be elicited by optogenetic activation of single retinorecipient neurons in the brain (Antinucci et al., 2019). How do RGC signals from the eye supply these circuits?

From optical recordings of RGC axon terminals in the brain, we have learnt that like in mammals (Baden et al., 2018), larval zebrafish RGCs are tuned to object size (Preuss et al., 2014) as well as orientation and motion direction (Nikolaou et al., 2012), each organised into specific layers and regions of the brain including the tectum, pretectum and thalamus (Hildebrand et al., 2017; Nikolaou et al., 2012; Robles et al., 2014; Semmelhack et al., 2014; Wulliman et al., 1996). However, our understanding of RGC structure and function in zebrafish remains far from complete.

First, zebrafish have a large field of view that lets them simultaneously survey the overhead sky and the riverbed beneath them (Bianco et al., 2011; Patterson et al., 2013; Zimmermann et al., 2018). These parts of visual space have vastly different behavioural relevance, as well as distinct spatial, temporal and spectral statistics (Baden et al., 2020; Engeszer et al., 2007; Nevala and Baden, 2019; Parichy, 2015; Zimmermann et al., 2018). For efficient coding (Attneave, 1954; Barlow, 1961; Simoncelli and Olshausen, 2001) zebrafish should therefore invest in different sets of functional RGC types to support different aspects of vision across their retinal surface. In agreement, both photoreceptor (Yoshimatsu et al., 2019) and retinal bipolar cell functions (Zimmermann et al., 2018) are asymmetrically distributed across the eye, and feature pronounced reorganisations in the *area temporalis* (dubbed Strike Zone, SZ (Zimmermann et al., 2018)), which is used for visual prey capture (Bianco et al., 2011; Mearns et al., 2019; Semmelhack et al., 2014; Yoshimatsu et al., 2019; Zimmermann et al., 2018). In contrast, data on structural and functional retinal anisotropies in zebrafish RGCs remains largely outstanding (but see (Robles et al., 2014)).

Second, characterising RGC functions by recording the signals of their axonal arborisations in the brain is limited by the fact they are densely packed, meaning that it is difficult to segment signals belonging to individual RGCs in population recordings (Nikolaou et al., 2012). In hand, RGC axon terminals are potentially subject to central presynaptic inputs (Ferguson and McFarlane, 2002; Ramdya and Engert, 2008; Sajovic and Levinthal, 1983), which leaves it unclear which aspects of their response properties first emerge in the eye, and which emerge only in the brain.

Third, most investigations into the function of zebrafish visual circuits have relied on long-wavelength light stimulation to limit interference with fluorescence imaging systems (Bollmann, 2019). However, like many surface-dwelling teleost fish (Baden and Osorio, 2019; Champ et al., 2016; Neumeyer, 1992), zebrafish have rich tetrachromatic colour vision (Krauss and Neumeyer, 2003; Meier et al., 2018; Orger and Baier, 2005), and spectrally diverse functions dominate both their outer (Klaassen et al., 2016) and inner retinal circuits (Connaughton and Nelson, 2010; Meier et al., 2018; Zimmermann et al., 2018). In fact, wavelength is strongly associated with specific behaviours in zebrafish. For example, the optomotor response is best driven by long wavelength light (Orger and Baier, 2005), while visual guided prey capture capitalises on short-wavelength vision (Yoshimatsu et al., 2019). However, if and how zebrafish visual behaviours build on signals from spectrally selective RGC circuits remains unknown.

To address these major gaps in knowledge, we systematically imaged light-driven signals from RGCs directly in the *in vivo* eye. By ‘bending’ the imaging scan-plane to follow the natural curvature of the live eye (Janiak et al., 2019), and synchronising the stimulation light with the scanner retrace (Baden et al., 2013; Euler et al., 2019; Franke et al., 2019; Zimmermann et al., 2018), we chart the *in vivo* functional diversity of larval zebrafish RGCs in time and wavelength across visual space.

We find that zebrafish RGCs support a broad range of both achromatic and chromatic functions and display a notable interdependence of temporal and spectral signal processing. Nearly half of all RGC processes display temporally complex forms of colour-opponency, including many that are emerge from a pervasive presence of slow blue-Off signals across the retina. The latter may be linked to a spectral segregation of short- and long-wavelength driven circuits with distinct behavioural relevance. Next, RGC functions and their organisation across the inner retina varied strongly with position in the eye, including a striking regional prominence of UV-sensitive sustained ON-circuits in the SZ. This part of the eye also had the highest density of RGCs, despite amacrine cell numbers (ACs) remaining approximately constant across the retinal surface. Finally, photoconversion experiments revealed an equally striking systematic morphological variation of individual RGCs in different parts of the eye: SZ RGCs tended to be small-field but diffusely stratified, while nasal RGCs tended to be more widefield but narrowly stratified. Together, our data strongly suggests that functionally and morphologically distinct types of RGCs occupy distinct parts of the zebrafish eye to serve distinct visual functions and point to the existence of a set of highly specialised sustained UV-On ‘prey-capture-RGCs’ in the SZ.

## RESULTS

### The density of RGCs, but not of ACs, is locally increased in the Strike Zone

Larval zebrafish use a highly asymmetrical retina to process distinct sets of visual features across different parts of visual space (Robles et al., 2014; Yoshimatsu et al., 2019; Zimmermann et al., 2018). In particular, they feature a pronounced acute zone (*area temporalis* (Schmitt and Dowling, 1999), dubbed “strike zone”, SZ; (Zimmermann et al., 2018)) which is used in conjunction with fixational eye movements for visual guided prey capture (Antinucci et al., 2019; Mearns et al., 2019; Semmelhack et al., 2014; Yoshimatsu et al., 2019). However, the anatomical distribution of RGCs across the eye has never been quantified. In larval zebrafish, the somata of RGCs reside exclusively in the ganglion cells layer (GCL), which also harbours displaced amacrine cells (dACs). To therefore establish the number and distribution of all RGCs across the intact 7 *dpf* eye we labelled all somata with DAPI and used the amacrine-cell specific promotor ptfa1 (Jusuf and Harris, 2009) to in addition label amacrine cells with mCherry. From here, we detected all DAPI labelled cells in the GCL (RGCs+dACs, n=5,750) as well as all mCherry-labelled cells in the GCL (dACs, n=765) and the inner nuclear layer (INL, ‘regular’ ACs, n=3,105), and projected each into a local distance-preserving 2D map (Fig. 1A,B_1-3_ (Zimmermann et al., 2018)). We then subtracted dACs from GCL cells to isolate a total of n = 4,985 RGCs (Fig. 1C_1_, see also (Robles et al., 2014)), and summed all ptf1a-positive cells to isolate a total of n=3,870 ACs (Fig. 1C_2_).

**Figure 1.**
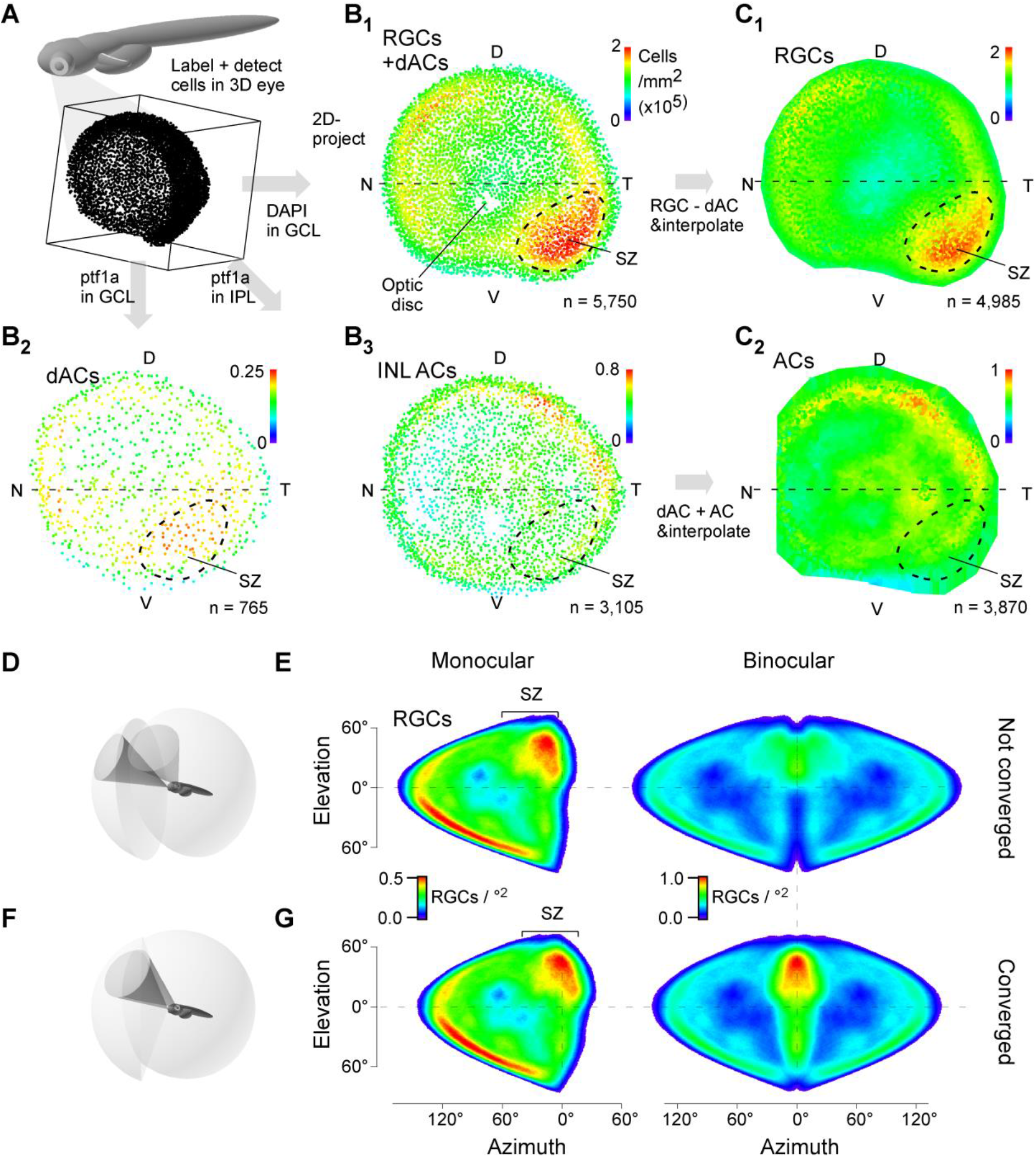
RGCs, but not ACs, are elevated in the strike zone. **A**, Schematic of larval zebrafish and enlarged 3D representation of GCL nuclei in the eye. **B_1-3_**, 2D projections of detected soma positions across the eye of all GCL cells based on a DAPI stain which includes all RGCs and all dACs (1) and selective isolation of amacrine cells in the GCL (dACs, 2) and INL (ACs, 3) based on ptf1a labelling. **C_1,2_**, Density maps of all RGCs (1) and ACs (2) computed from cell counts in (B). D, Dorsal; N, Nasal; V, Ventral; T, Temporal, SZ, Strike zone. **D-G**, 3D schematics (D, F) and projections of RGC densities (E, G) into monocular and binocular visual space during ‘rest’ (D, E, eyes not converged) and ‘hunting’ (F, G, eye converged).

As predicted from work on photoreceptors (Yoshimatsu et al., 2019; Zimmermann et al., 2018), the density of RGCs was elevated in the SZ (Figs. 1C_1_). In contrast, ACs were distributed approximately homogeneously. As a result, the SZ had a ~50% reduced AC:RGC ratio in comparison to the rest of the retina average of ~3 ACs per 4 RGCs. If this regional relative reduction in AC numbers also results in a relative reduction amongst inhibitory synapses remains untested. Conceptually reminiscent, a reduced inhibitory tone in the primate fovea is thought to aid signal-to-noise in low-convergence circuits (Baden et al., 2020; Sinha et al., 2017). In view of their need to process the signal from small numbers of cones for visual prey detection (Yoshimatsu et al., 2019), larval zebrafish RGCs might also be expected to benefit from a low-inhibition arrangement in their SZ.

When projected into binocular visual space, with eyes in their non-converged resting position, the two eyes’ SZs were aligned with the upper-frontal visual field, just ipsilaterally off-centre (Fig. 1D,E). The same regions in visual space are most effective in eliciting prey-detection behaviour (Bianco et al., 2011; Mearns et al., 2020). Upon eye convergence, the two eyes’ acute zones superimposed for binocular high-acuity vision, which likely aids estimating the relative 3D location of potential prey and ultimately trigger prey-capture behaviour (Fig. 1F,G). Nevertheless, due to the small size of the eye the larval zebrafish SZ ‘only’ comprises a total of ~400 RGCs per eye, suggesting that very low numbers of individual RGCs that both belong to specific types and are situated in the SZ can be tasked with sending critical information about the presence and position of prey to the brain. Conceptually in agreement, already single retinorecipient pre-tectal neurons are sufficient to elicit prey-capture-like behaviour upon optogenetic stimulation (Antinucci et al., 2019). What are the visual response properties of SZ-RGCs, and how do they compare to the functions of RGCs responsible for conveying information about the remainder of visual space? To address this question, we next turned to 2-photon imaging of RGCs in the live eye (Fig. 2).

**Figure 2.**
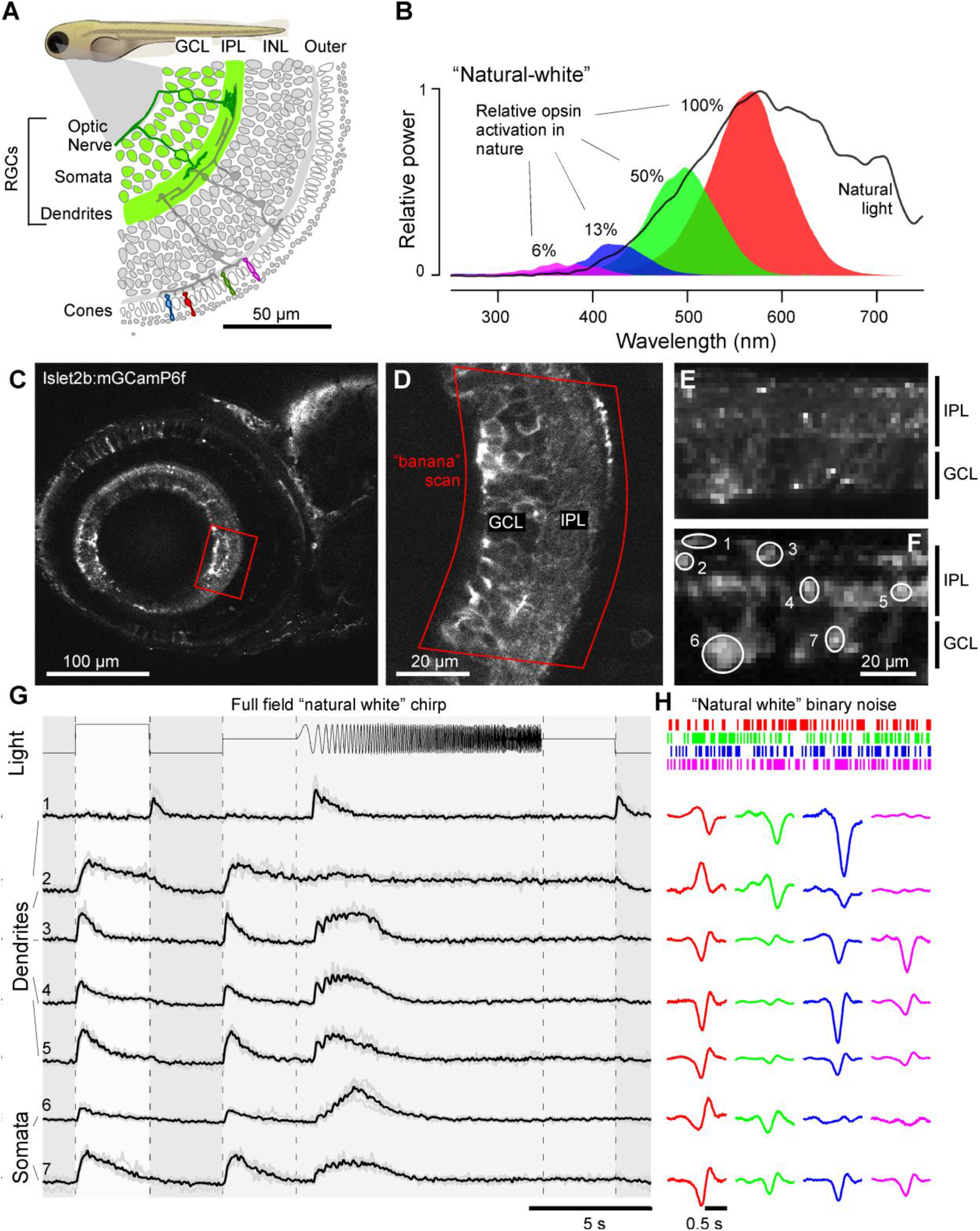
Recording from RGC dendrites and somata in vivo. **A**, Schematic of Islet2b:mGCaMP6f expression in RGCs (green) across a section of the larval zebrafish eye, with somata in the ganglion cell layer (GCL) and dendrites in the inner plexiform layer (IPL), see also Supplementary Fig. S1. INL Inner nuclear layer. **B**, Average spectrum of natural daylight measured in the zebrafish natural habitat from the fish’s point of view along the underwater horizon (solid line). Convolution of the zebrafish’s four cone action spectra with this average spectrum (shadings) was used to estimate the relative power each cone surveys in nature, normalised to red cones (100%). Stimulation LED powers were relatively adjusted accordingly (“natural white”). **C,D**, GCaMP6f expression under 2-photon surveyed across the entire eye’s sagittal plane (C) and zoom in to the strike zone as indicated (D). Within the zoomed field of view, a curved scan path was defined (“banana scan”) to follow the curved GCL and IPL for activity recordings (E) which effectively ‘straightened’ the natural curvature of the eye. **E**, Example activity scan with RGC dendrites occupying the top part of the scan in the IPL, and somata occupying the bottom part in the GCL as indicated (E, top) and correlation projection (Franke et al., 2017) of activity following white noise stimulation highlighting responding regions in the scan alongside example regions of interest (ROIs, E, bottom) (see also Supplementary Video S1) **G**, Mean (black) and individual repeats (grey) example responses of ROIs from (E) to full field stimulation as indicated. **H**, As (G), now showing linear kernels to red, green, blue and UV components recovered from natural white noise stimulation (Methods). Note that several ROIs display a robust UV-component despite the ~20-fold attenuated stimulation power in this band relative to red (B). See also Supplementary Fig. S2.

### Highly diverse light-driven responses of RGCs in the live eye

To record light-driven activity from RGC processes in the eye we expressed the membrane-tagged variant of GCaMP6f (mGCaMP6f) under the RGC-associated promoter Islet2b (Janiak et al., 2019; Thisse et al., 2004). This reliably labelled the somata and dendrites of most RGCs (Fig. 2A, Supplementary Fig. S1, Supplementary Discussion). For stimulation, we presented full-field light modulated in time and wavelength based on four LEDs that were spectrally aligned with the sensitivity peaks of the zebrafish’s four cone opsins (R, G, B, UV) (Zimmermann et al., 2018). The power of each LED was adjusted to follow the relative power distribution across wavelength of daytime light in the zebrafish natural habitat (Nevala and Baden, 2019; Zimmermann et al., 2018) to yield a “natural white”: red (100%), green (50%), blue (13%) and UV (6%) (Fig. 2B). This adjustment was to ensure that recorded RGC’s spectral responses would be maximally-informative about their likely performance in a natural setting. For example, our previous study of spectral tuning of zebrafish BCs revealed a profound UV-dominance in the upper-frontal visual field when probed with spectrally flat light, which left it unclear how much of this dominance can be explained by a possible eye-wide increase in UV-sensitivity compared to other spectral cones. Remarkably, although high-UV power stimulation clearly affected the overall waveforms of RGC responses to noise stimulation, over prolonged stimulation this resulted in no significant difference in the amplitudes and distributions of spectral receptive fields (Supplementary Fig. S2A-E).

Animals were imaged under 2-photon at 6-8 *dpf*. All recordings were performed in the eye’s sagittal plane (Fig. 2C). In each case, after zooming in on given region of the eye, we ‘bent’ the scan to follow the curvature of the eye (Fig. 1D. ‘banana scan’, Methods). This allowed recording the entire width of the IPL and GCL without sampling adjacent dead-space (Fig. 1E). Conveniently, this strategy effectively ‘un-bent’ the natural curvature of the eye thus facilitating analysis (Methods): An example 15.6 Hz recording at 64×32 pixel resolution comprised a ‘straightened’ IPL in the upper part of the image, and the GCL in the lower part (Fig. 2E,F, Supplementary Video S1). Together, this allowed sampling both RGC dendrites, which integrate inputs from BCs and ACs (IPL) (Baden et al., 2018; Masland, 2012; Ran et al., 2019), as well as from RGC somata, whose activity is expected to largely reflect the spiking activity for transmission to the brain (GCL) (Baden et al., 2016). Throughout, we present data recorded from these distinct structures together, with dendrites plotted on top and somata plotted on an inverted y-axis below (e.g. Fig. 3A-C). By *en-large*, and with exceptions noted in the relevant sections below, the types and distributions of dendritic and somatic functions tended to be largely in line with each other.

**Figure 3.**
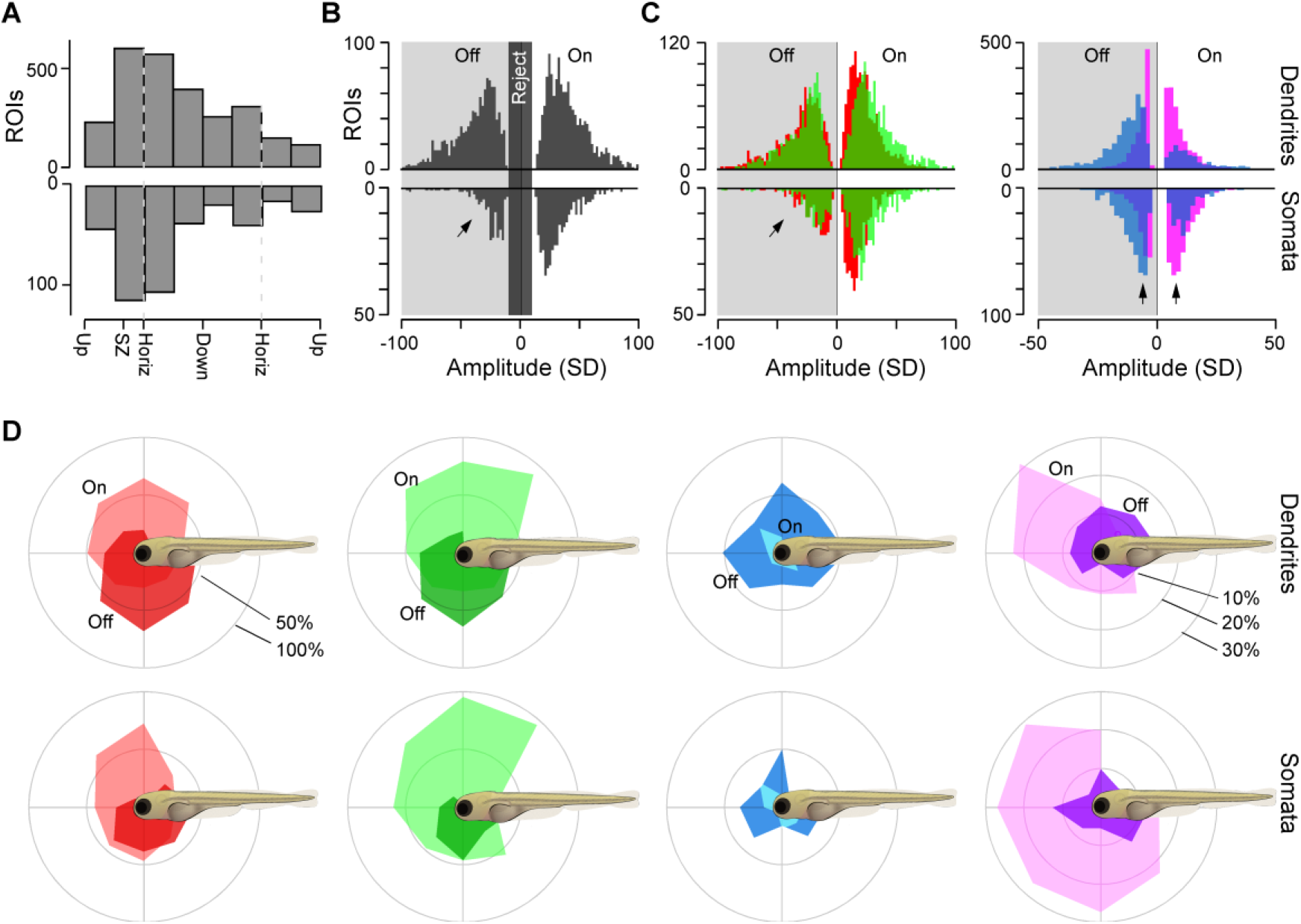
Overview of dendritic and somatic RGC functions across the eye. **A**, Number of dendritic (top) and somatic (bottom, y-flipped) ROIs recorded across different positions in the eye. The relative abundance of SZ-ROIs is in line with the increased RGC numbers in this part of the eye (cf. Fig. 1). **B, C**, Kernel amplitudes of all dendritic (top) and somatic (bottom, y-flipped) ROIs. Shown for the maximal amplitude kernel of each ROI irrespective of colour (B) and separated by colour (C) with red/green (left) and blue/UV (right) plotted separately for clarity. Arrowheads emphasise a relative reduction in red/green OFF responses at the level of somata (left) and dominance for Off- and On-responses across the entire dataset for blue and UV-kernels, respectively (right). **D**, Prominence of different colour and polarity responses (from C) amongst dendrites (top row) and somata (bottom row), plotted across visual space. In each case, all kernels that exceeded a minimum amplitude of 10 SDs were included. Scalebars in percent of dendritic/somatic ROIs that were recorded in a given section of the eye such that the percentages of On, Off and non-responding (<10 SD) add to 100%.

For each scan, we presented two stimuli: A ‘natural-white’ time varying chirp stimulus (Baden et al., 2016) to assess RGCs’ achromatic response properties, and a 6.4 Hz natural-power spectrum tetrachromatic binary noise stimulus to probe their spectral tuning (Zimmermann et al., 2018). Reverse correlation of each ROIs’ response to this stimulus allowed computing four linear kernels, one for each stimulated waveband (Methods).

In an example recording, a selection of regions of interest (ROIs) revealed a rich diversity of response properties across both RGC dendrites and somata (Fig. 2G,H). For example, dendritic ROI 1 was a blue-biased transient Off-process, while immediately adjacent ROI 2 was a “red vs. green/blue” colour opponent sustained On-process. Similarly, also different RGC somata responded in diverse manners: ROI 6 exhibited a red-dominated transient On response with a band-pass response in the frequency domain, while ROI 7 was a largely achromatic On cell. We next systematically recorded RGC responses to these stimuli across different positions in the eye.

### RGCs’ polarities and spectral response properties vary across visual space

In total, we recorded 72 such fields of view (n = 17 fish), and automatically placed ROIs on functionally homogeneous processes based on local response correlation during the tetrachromatic noise stimulus (Franke et al., 2017) (Supplementary Fig. S2F-H, Methods). Each ROI was categorised as either dendrite or soma based on its vertical position in the scan. ROIs from the SZ were relatively overrepresented (Fig. 3A), in line with the relative RGC-density elevation (Fig. 1C_1_) and IPL thickening (Zimmermann et al., 2018) in this part of the eye.

From here, low-amplitude ROIs were discarded (Methods) and thereafter classed as either dominant “On” or “Off” based on the dominant sign of their largest amplitude kernel (Fig. 3B, Methods). Under this set of criteria, dendritic ROIs were approximately evenly (54:46 On:Off) divided into the On and Off groups (n = 1,461 On, 1,255 Off), while somata comprised relatively more On ROIs (66:34 On:Off; n = 388 On, 198 Off; Discussion). Similarly, when considering only red or green kernels individually, On strongly dominated at the level of somata (red: 65% On: n=378 On, 208 Off; green 85% On: n = 416 On, n = 70 Off) but not dendrites (red: 47% On: n = 1,291 On, 1,452 Off; green: 43% On: n = 1,164 On, 1,552 Off) (Fig. 3C). In contrast, both at the level of somata and dendrites, blue kernels were strongly Off-biased (somata: 67% Off: n = 196 On, 390 Off; dendrites: 73% Off: n = 732 On, 1,984 Off), while UV somatic but not dendritic kernels were On-biased (somata: 64% On: n = 378 On, 211 Off; dendrites: 44% On: n = 1,192 On, 1,542 Off).

Next, we computed how On and Off-type responses in each waveband varied across the eye and thus across corresponding position in visual space. This revealed that On- and Off-processes were generally biased to the upper and lower visual fields, respectively, in line with our previous findings from bipolar cells (Zimmermann et al., 2018). However, blue-Off RGC processes dominated over blue-On processes throughout visual space. Finally, both On and Off UV-processes mostly surveyed the upper visual field. However, UV-On processes were strongly biased to the frontal-upper visual field, likely to support visually guided prey capture of UV-bright plankton (Yoshimatsu et al., 2019), while UV-Off processes approximately evenly surveyed upper visual space without any obvious bias for the frontal visual field. The latter may support UV-dark silhouette detection (Cronin and Bok, 2016; Yoshimatsu et al., 2019) above the fish, for example for predator avoidance. How are these broad organisational differences established within the layers of the IPL?

### RGC dendrites simultaneously encode contrast, time and colour

To determine the dominant functional properties of RGC processes in different parts of the eye, we mapped each dendritic ROI to a bin within an “Eye-IPL map”. In this representation, the x-coordinate denotes position across the eye (dorsal, nasal etc.) while the y-coordinate represents IPL depth (Fig. 4A,B). We then computed each Eye-IPL bin’s mean light response to the chirp stimulus and projected its time axis into the third dimension to yield an array linking eye position (x), IPL position (y) and time (z) (Fig. 4C). In this representation, the spatially resolved mean response of all RGC dendrites could be visualised as a movie (Supplementary Videos S2,3). Alternatively, the mean RGC response in a given region in the eye could be displayed as a trace over time (Fig. 4D) or individual time-points could be displayed as images over Eye-IPL-space (Fig. 4E). This analysis revealed that polarity, transience and frequency tuning of RGC dendrites all varied systematically across the eye.

**Figure 4.**
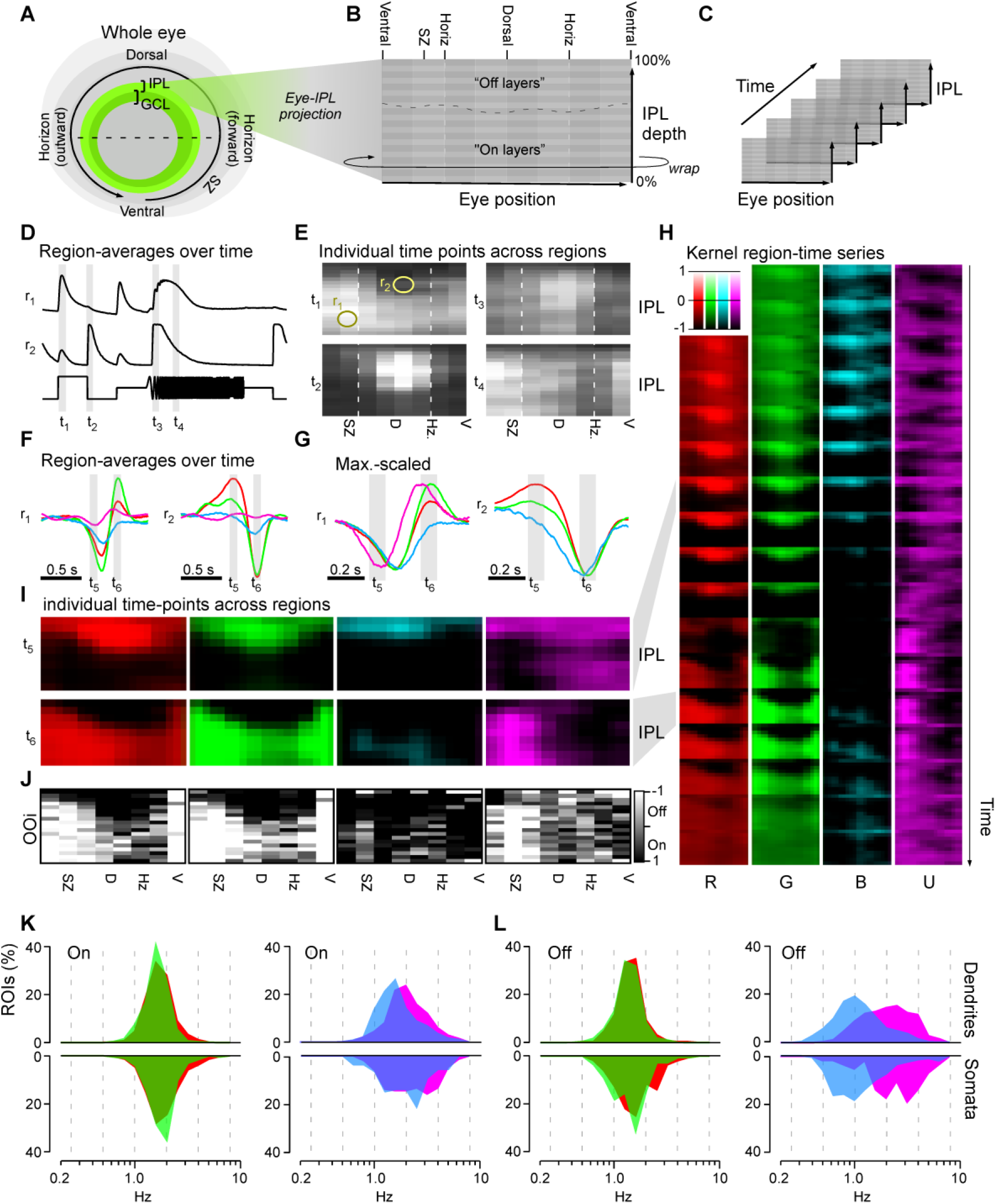
Major functional response trends across the eye and IPL. **A-C**, Schematic illustrating how dendritic ROIs from different parts of the eye and IPL depth (A) were mapped into a 2-dimensional “Eye-IPL” map (B), which can then also be analysed over time (C). Note that this involved ‘cutting’ the circular range of eye positions, such that the ventral retina is represented at either edge along the 2-projections’ x-axis. **D,E**, Example snapshots of mean responses to chirp stimulation (cf. Fig. 2G) mapped into an eye-IPL map as schematised above (A-C). Data can be plotted as time traces for a given region of the eye and IPL (D, r_1,2_ as indicated in E), or alternatively as a time-frozen snapshot of activity across the eye and IPL at different points in time (E, t_1-4_ as indicated in D). See also Supplementary Videos S2,3. **F-I**, as (D,E), but instead showing mean kernels across the four spectral wavebands, where F,G are mean and max-scaled mean kernels for Eye-IPL regions r_1,2_ (as in E), respectively. (H) shows the evolution of max-scaled kernel Eye-IPL maps over consecutive timepoints as indicated, with (I) showing an enlarged version of the same plot for timepoints t_5,6_ (as indicated in F, G and H) including colour-scalebar which also apply to (I), where 0 equates to the baseline of each bin’s kernel, and 1/-1 to their respective maximum or minimum (cf. G). See also Supplementary Video S4. **J**, Projection of an On-Off index (OOi, Methods) in the four wavebands (as in I) into an Eye-IPL map, with yellow and blue shades indicating an overall On- and Off-bias, respectively (see also Fig. 3D). **K**, Distribution of central frequencies (Methods) of dendritic (top) and somatic (bottom, inverted y-axis) kernels in the four wavebands, separated into On (K) and Off (L) kernels. For clarity, red/green (left) and blue/UV (right) data is plotted separately in each case. Wilcoxon Rank Sum test, 1 tailed with correction for multiple comparisons for all pairwise comparisons between same polarity distributions of spectral centroids. Dendrites: all p<0.001 except R_Off_ vs. G_Off_ (p=0.0011) and G_On_ vs. B_On_ (p=0.69). Somata: all p<0.001 except R_On_ vs. U_On_ (p=0.00101), R_Off_ vs. G_Off_ (p=0.033), G_On_ vs. B_On_ (p=0.045), B_On_ vs. U_On_ (p=0.064), R_On_ vs. B_On_ (p=0.25) and R_On_ vs. G_On_ (p=0.57).

For example, a region in the SZ’s On-layer (region 1, r_1_) on average responded to the onset of a flash of light and exhibited broad frequency tuning during temporal flicker (Fig. 4D, top). In contrast, a region within the dorsal eye’s Off-layer (r_2_) on average exhibited an Off-dominated transient On-Off-response and low-pass tuning to temporal flicker (Fig. 4D, bottom). Vice versa, inspection of individual time points (t_1-4_) revealed a strong asymmetry in the distribution of these response properties across both the IPL (y) and the eye (x) (Fig. 4E). For example, rather than forming two straight horizontal bands of On- and Off responses, the position of the On-Off boundary varied strongly across the eye (t_1,2_ in Fig. 4E). Off responses dominated much of the IPL in the dorsal retina but were compressed to a mere ~10% of IPL width in the ventral part. Also the mean temporal frequency preference varied across the eye: The dorsal-most retina exhibited the most low-pass tuning to temporal flicker, while increasingly ventral regions progressively switched to bandpass tuning (t_3,4_ in Fig. 4E, best seen in Supplementary Video S3). In this achromatic regime, different parts of the eye therefore on average encoded the polarity and speed of visual stimuli in very different ways. We next asked how these properties were linked to the zebrafish’s four spectral input channels.

For this we mapped the spectral kernels into the same reference frame. This yielded four kernel-movies, one each for red, green, blue and UV stimulation (Supplementary Video S4). We first compared the temporal profiles across the same regions r_1_ and r_2_ as before. In line with the achromatic chirp response (Fig. 4D), r_1_ was dominated by On-kernels, while r_2_ was dominated by Off-kernels (Fig. 4F,G). However, in each case time-courses varied greatly between spectral bands. For example, r_1_ exhibited a biphasic UV-On-kernel, temporally offset biphasic On-kernels in red and green, and a monophasic blue Off-kernel. Similarly, r_2_ exhibited three distinct temporal profiles across red (biphasic), green (weakly biphasic) and blue (monophasic). Accordingly, spectral information was not only encoded through variations in gain and polarity of RGC responses but was in addition mixed with temporal information – reminiscent of a previous similar observation in adult zebrafish amacrine cells (Torvund et al., 2017) as well as in the locust visual system (Osorio, 1986, 1987).

To more systematically explore how wavelength- and time-information interplay across different regions of the eye and IPL, we plotted the kernel-movies as a time series (Fig. 4H), and specifically highlighted the two time points that aligned with the peaks of most kernels’ On- and Off-lobes (t_6_ and t_5_, respectively, Fig. 4I). In this representation, the red and green kernel-maps were highly reminiscent of the achromatic On (t_1_) and Off (t_2_) response profiles during chirp stimulation (Fig. 4I, cf. t_1,2_ in Fig. 4E). In contrast, blue kernels consistently lacked a dominant On-lobe (Fig. 4I, blue, bottom), in line with their overall Off-dominance (cf. Figs. 3C,D). Finally, UV-kernels were different still: In the SZ, their IPL-depth profile approximately resembled red/green kernels (Fig. 4I, magenta), while in the remainder of the eye, much of the On-band seen in red/green instead transitioned into a secondary UV-Off-band (Fig. 4I, magenta, top). To quantify the differences in the distribution of On- and Off-signals, we computed an On-Off index (OOi, Methods). OOis of 1 and −1 denote regions exclusively comprised of On and Off kernels, respectively, while an OOi of zero denotes an equal proportion of On and Off kernels. The resultant OOi maps confirmed the differential distributions of On and Off signals seen across in the individual kernel-maps (Fig. 4J).

Next, we considered the temporal domain. As across Eye-IPL space, red and green maps continued to resemble each other (Fig. 4H). In contrast, the blue-map was consistently slowed across the entire eye, while the UV-map exhibited a complex temporal behaviour that in addition strongly differed between the SZ and the remainder of the eye (Fig. 4H, best seen in Supplementary Video S4). These broad differences were also evident from the kernels’ central frequencies (here: each kernel’s spectral centroid determined from Fourier transform) irrespective of eye-position (Fig. 4K,L). Red and green kernels exhibited a narrow range of intermediate central-frequencies, while blue kernels were systematically slowed down and UV-kernels were systematically sped up. These differences were particularly pronounced for Off- (Fig. 4L) compared to On-kernels (Fig. 4K).

Together, this functional overview strongly suggests that (1) information received across the four different wavebands of light are used in distinct ways to support vision, and (2) their use varies strongly across position in the visual field (Baden et al., 2020) (see Discussion). To further explore how spectral information might serve zebrafish vision at the level of the retina’s output we next assessed RGC responses for spectral opponency.

### An abundance of temporally complex colour opponent RGCs

Colour opponency in the retinal output is often taken as a hallmark of circuits for colour vision (Baden and Osorio, 2019). Previous behavioural and physiological experiments have clearly demonstrated tetrachromatic colour vision including colour constancy in species of cyprinids (Dörr and Neumeyer, 2000; Krauss and Neumeyer, 2003; Neumeyer, 1992) including key aspects in zebrafish (Meier et al., 2018). This implies that there should be at least three spectrally distinct mechanisms of opponency in the visual system. We therefore wondered how their RGCs might support their colour vision.

When combining the signal from multiple cone pathways for output to the brain, the number of possible wiring combinations is given by the number of possible wiring states (i.e. 3: On, Off, no connection) raised to the power of the number of cone types (i.e. 4). Accordingly, the zebrafish’s four cone types could be wired in a total of 3^4^ = 81 combinations. Of these, 50 are colour-opponent, 30 are non-opponent (15 On + 15 Off), and one represents the case where none of the four cones is functionally connected. We assessed how zebrafish RGCs span this combinatorial space and ranked the results based on the number of allocated dendritic ROIs in each wiring group (Fig. 5). This revealed that most ROIs fell into a small subset of groups with relatively simple functional wiring motifs. Amongst dendrites, the two most common combinations were RGB_Off_ and RG_Off_ (Fig. 5A, top, dark grey and Fig. 5B, groups 1,2). These non-opponent Off groups were followed by one colour opponent group (RG_On_-B_Off_, brown/orange; group 3) and then two non-opponent On groups (RG_On_ and RGB_On_, light-grey; groups 4,5). Together, these first five groups made up 42% of all dendritic ROIs. However, subsequent groups were more diverse and largely comprised of colour opponent categories to make up a total of 47% colour opponent ROIs amongst dendrites (e.g. Fig. 5B groups 6, 7, 9, 10). Of these, most (75%) opponent computations had a single zero crossing in wavelength: R/G (30%), G/B (31%), B/U (8%), G/U (4%) and RU (2%), respectively (e.g. Fig. 5B groups 3, 6, 10). The remaining 25% of opponent ROIs described diverse complex opponencies (e.g. Fig. 5B group 7, 9). A similar distribution of functions was found for somata (51% non-opponent, 49% opponent - of which 67% and 33% exhibited simple and complex opponencies, respectively, Fig 5A, bottom), with the notable exception of a drop in the first two Off groups (cf. Fig. 3).

**Figure 5.**
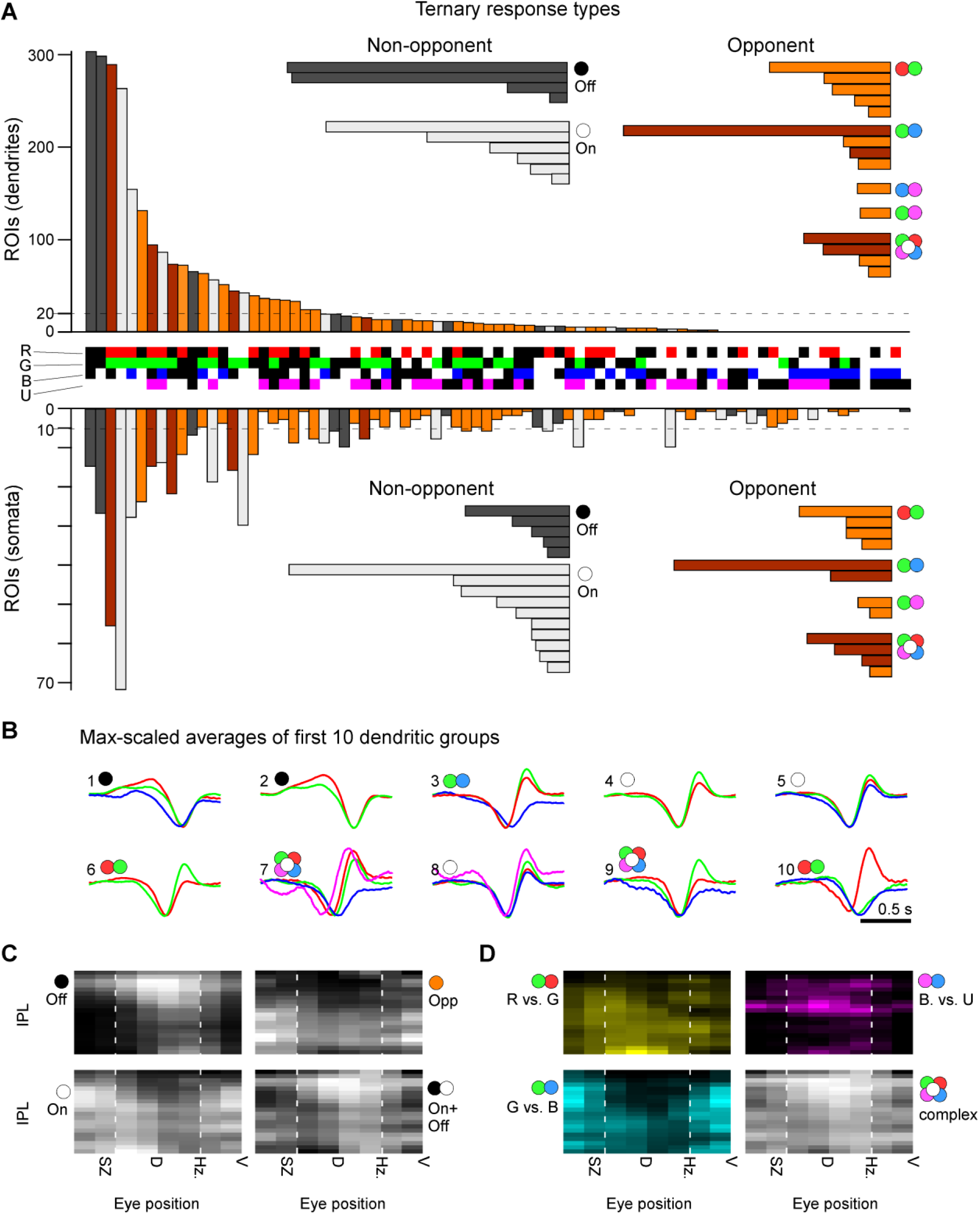
Diverse colour opponencies in RGCs. **A**, Each dendritic (top) and somatic (bottom, inverted y-axis) ROIs that passed a minimum response criterion (Methods) was allocated to a single bin in a ternary classification scheme according to the relative polarities of their four spectral kernels (3 response states On, Off, no response) raised to the power of 4 spectral channels (red, green, blue, UV): 3^4^ = 81 possible combinations. The central row between the bar graphs indicates each bin’s spectral profile: “On” (red, green, blue, UV), “Off” (black in the respective row) and no response (white in the respective row). For example, the leftmost group, which comprised the highest number of dendritic ROIs, corresponds to ROIs displaying Off kernels in red, green and blue, with UV showing no response. The bar graphs are colour coded as follows: dark grey (non-opponent Off), light grey: (non-opponent On), Orange/brown (opponent). Brown bins indicate opponent bins that are only classified as opponent because they comprise a Blue-Off component (see main text). The dotted horizontal lines denote the threshold of minimal ROI numbers included in a given bin to be included in the horizontally oriented summary insets. Colour circles next to these summary insets denote each group’s main spectral computation, with two-colour symbols denoting “simple” opponencies (single zero crossing) while the “flower” symbol denotes complex opponencies (>1 zero crossing). **B**, maximum-amplitude scaled average kernels of the ten most abundant spectral classes amongst dendrites in (A). **C, D**, dendritic groups from (A) summarised according to their position in an Eye-IPL map (cf. Fig. 4). (C) summarises major groups: Off (left, top) and On non-opponent (left, bottom), opponent (right, top), and On+Off non-opponent (right, bottom). (D), as (C), with opponent groups divided into their specific spectral computations as indicated. Note that most specific functions in (C,D) are restricted to specific regions of the eye and IPL. For example, G/B simple opponent computations occur mostly in the ON-layers of the the ventral retina that survey the world above the fish (D, bottom left).

As before (cf. Fig. 4) these diverse functional groups of non-opponent and opponent RGC processes also distributed asymmetrically across the eye and IPL depth (Fig. 5C,D). While colour-opponent RGC responses could be found across the eye, different types of colour-opponent computations dominated different parts of the IPL and visual field (Fig. 5D). For example, B/U opponent responses were mostly restricted to a narrow band in dorsal eye’s Off-layer, while in contrast G/B computations were mostly restricted to the ventral retina. R/G computations were comparatively more broadly distributed, but like B/U computations exhibited a preference for the dorsal retina. Notably, these distributions of spectral computations across the eye are only partially overlapping with colour-opponent responses previously observed at the level of bipolar cells (Zimmermann et al., 2018). In particular, unlike in colour-opponency in BCs, which was generally in the form of opposite sign but otherwise but temporally symmetrical spectral kernels, colour-opponency in RGCs was consistently mixed with temporal differences amongst opposite sign kernels (e.g. Fig. 5B, groups 3, 6, 7, 9, 10). Together, this hints at the presence of extensive further processing of spectral information beyond BCs, possibly involving spectrally diverse ACs (Torvund et al., 2017). We next wondered how this spectral information in the retinal output might map onto functional RGC types.

### UV-sensitive RGCs for prey capture in the strike zone?

While sorting RGCs based on their relative polarities to different wavelength light is instructive to capture details in the distribution of spectral computations (Fig. 5), it misses key temporal and amplitude information. To therefore more comprehensively identify the major functional RGC types of the larval zebrafish eye, we turned to clustering of RGCs’ full temporo-chromatic response profiles (Methods). This allocated dendritic ROIs into 18 functional clusters, of which 15 (C_1-15_) that contained a minimum of 10 members were kept for further analysis. These included largely achromatic On- (C_1,10,12_) and Off- (C_11, 13-15_) clusters as well as diverse clusters that displayed a mixture of spectral and temporal response properties (C_2-9_) (Fig. 6). However, unlike after sorting by opponency alone (Fig. 5), when clustered by this wider range or response properties, opponency was a less obvious feature across RGC groups. Moreover, where opponency was present, it was often primarily driven by the sluggish B_Off_ component opposing non-blue On-kernels (Fig. 6A,D-F, see also Fig. 6H). In fact, only four clusters did not exhibit an obvious sluggish B_Off_ response: C_3_, which did not respond to short wavelength stimulation at all, as well as the three achromatic On clusters (C_1,10,12_). This suggests that despite the pervasive presence of this subtractive B_Off_ signal throughout the eye, it is nevertheless used in a functional RGC-type specific manner.

**Figure 6.**
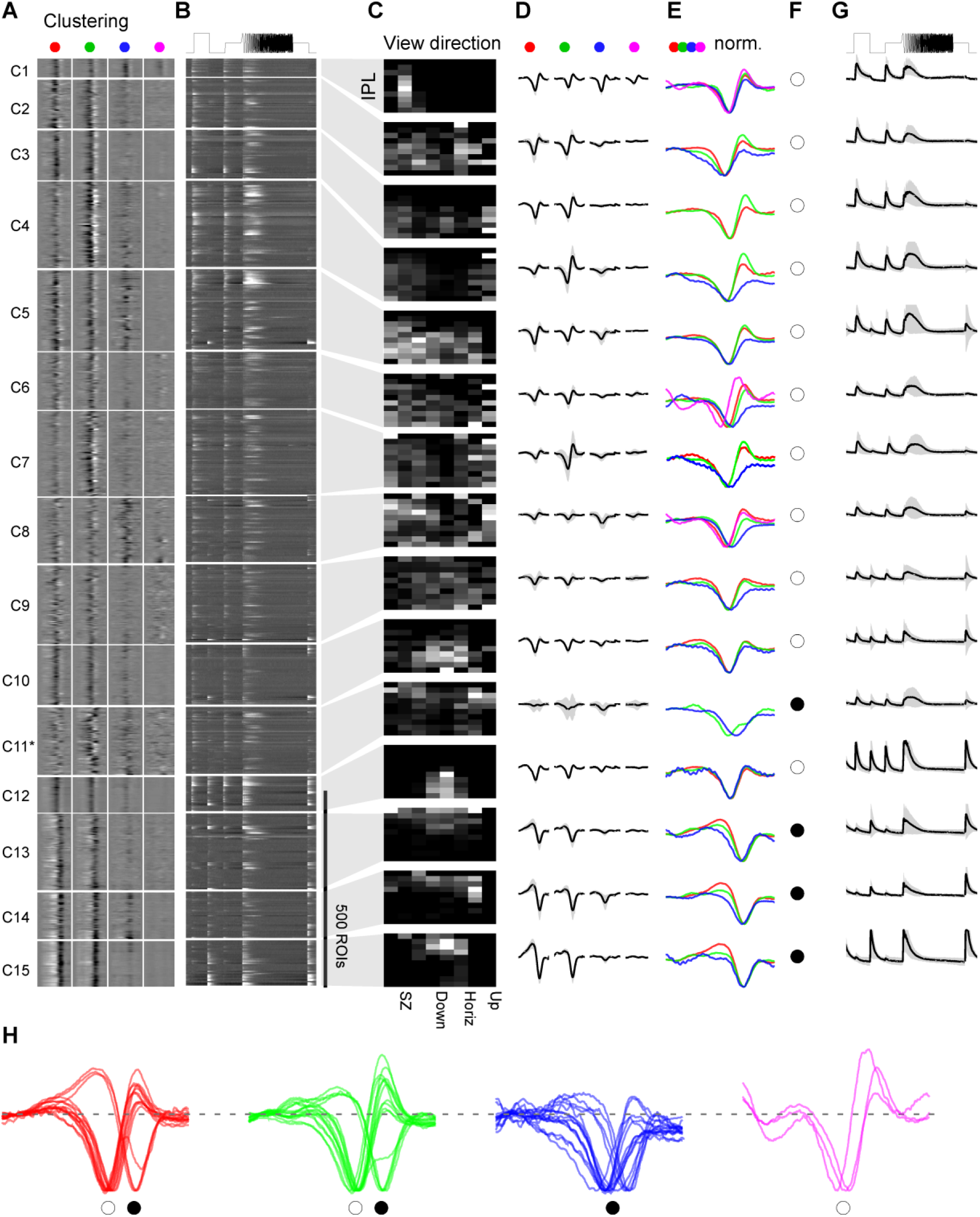
Functional clustering of dendritic ROIs. **A-G**, Dendritic ROIs from across the entire eye were clustered based on their four spectral kernels (Methods) to yield a total of n = 16 functional clusters that comprised a minimum of 10 ROIs. Note that Cluster C_12_* comprised a mixture of responses. Shown are heatmaps of red, green, blue, and UV kernels (A, from left to right, respectively) and associated mean chirp response (B), with each entry showing a single ROI, followed by each cluster’s Eye-IPL projection (C), each mean kernel (D), mx-scaled kernels superimposed (E), a spectral categorisation label (F) and the mean chirp response (G). Error shadings in s.d.. For clarity, low amplitude mean kernels were omitted from column (E). Spectral categorisation labels in (F) indicate the dominant polarity of each cluster: On (white) or Off (black). Note that C_11_* captured diverse kernels and may comprise a variety of low-n functional RGC types. For corresponding data on somata, see Supplementary Fig S3. **H**, as (E), but with all spectral kernels in each waveband enlarged and superimposed. Note kinetic similarities across most red and green kernels, and near complete absence of positive deflections in blue kernels. Greyscale colour maps (A-C) were linearly equalised by hand to maximise subjective discriminability of the full response range across the population of all recordings in a dataset. Lighter greys indicate higher activity / kernel amplitudes.

Next, most clusters exhibited a strong regional bias to either the upper or lower visual field (Fig. 6C). This provides further evidence that, like bipolar cells (Zimmermann et al., 2018), functional RGC types are asymmetrically distributed across the eye. Of these, C_1_ stood out in that it was the only one that responded strongly to UV-stimulation (Fig. 6A, D, E) despite the ~17-fold reduced signal power in our UV-stimulation light compared to red to match natural light (cf. Fig. 2B). This sustained On-cluster (Fig. 6G) remained tightly restricted to a single regional bin, which corresponded to the SZ (Fig. 6C). A functionally very similar and SZ-restricted cluster also featured amongst somatal ROIs (C_2_ in Supplementary Fig. S3). In view of the strong regionalisation of behavioural responses to prey-like stimuli (Bianco et al., 2011; Gebhardt et al., 2019; Mearns et al., 2019), and the strong facilitatory effect of UV light in prey-capture performance (Yoshimatsu et al., 2019), this strongly suggested that dendritic C_1_ and somatal C_2_ comprised the subset of RGCs responsible for visual guided prey-capture in larval zebrafish (Antinucci et al., 2019; Semmelhack et al., 2014; Yoshimatsu et al., 2019; Zimmermann et al., 2018). Nevertheless, likely in part due to their extreme regional restriction, in each case these putative “prey-capture” clusters only made up a tiny fraction of ROIs in this dataset, (3-5% amongst dendrites and somata, respectively). To therefore gain more in-depth information on the retina’s output from this part of the eye, we recorded and analysed a second functional dataset, but this time restricted all recordings to the strike zone (Figs. 7,8, Supplementary Fig. S4).

**Figure 7.**
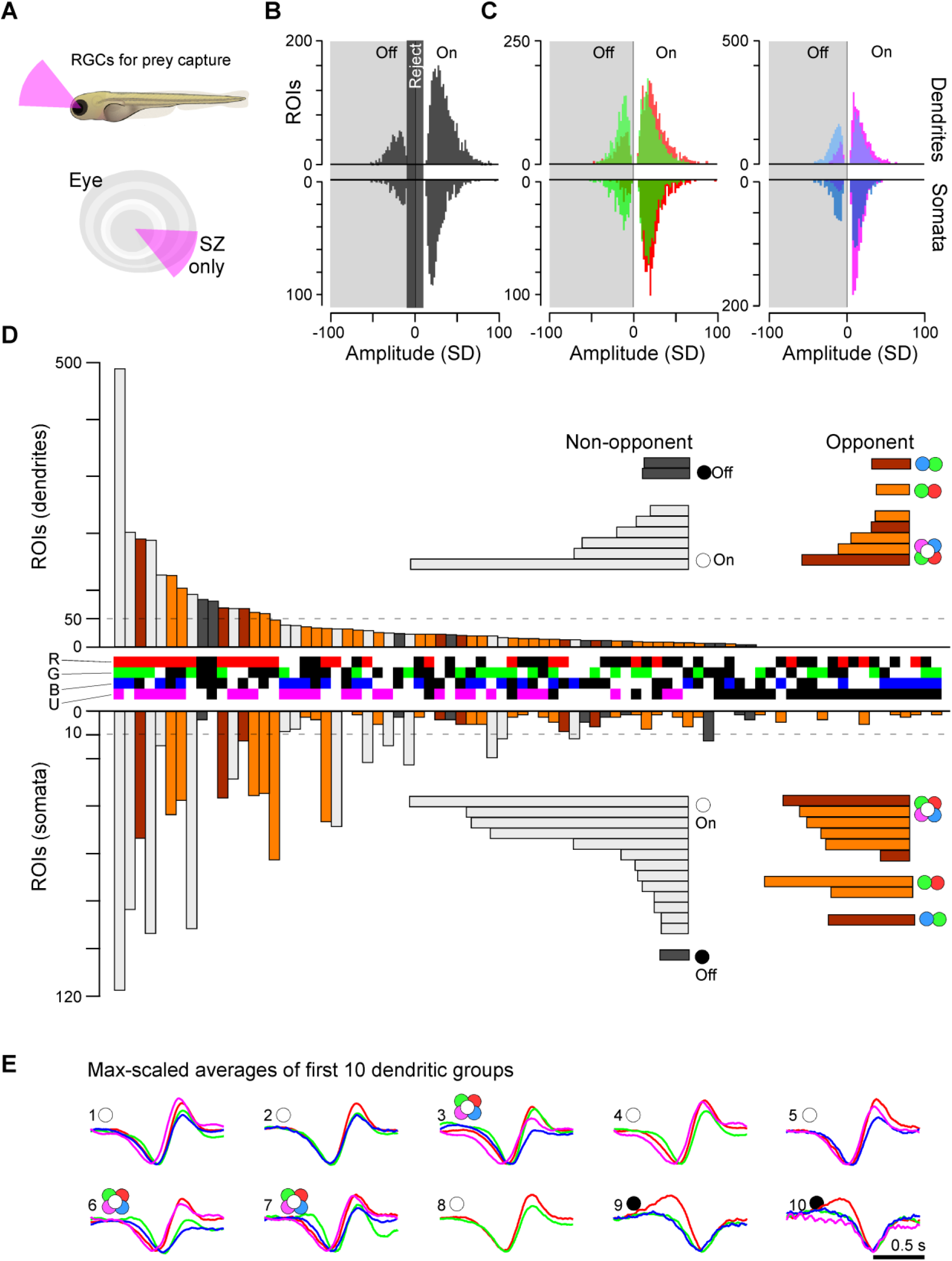
A closer look at function in the strike zone. **A**, A second series of RGC imaging experiments as shown in Figs. 2–6 was performed, this time exclusively recording from the strike zone (SZ), which surveys visual space above the frontal horizon. **B, C**, Overview of dominant On- and Off-responses (B) and by colour (C) amongst dendrites (top) and somata (bottom) for the SZ (for details cf. Fig. 3B,C). **D**, Ternary spectral classification of SZ dataset (for details cf. Fig. 5). Overall, note the striking On-dominance and increased presence of UV-responses in this dataset. **E**, maximum-amplitude scaled average kernels of the ten most abundant spectral classes amongst dendrites in (D).

**Figure 8.**
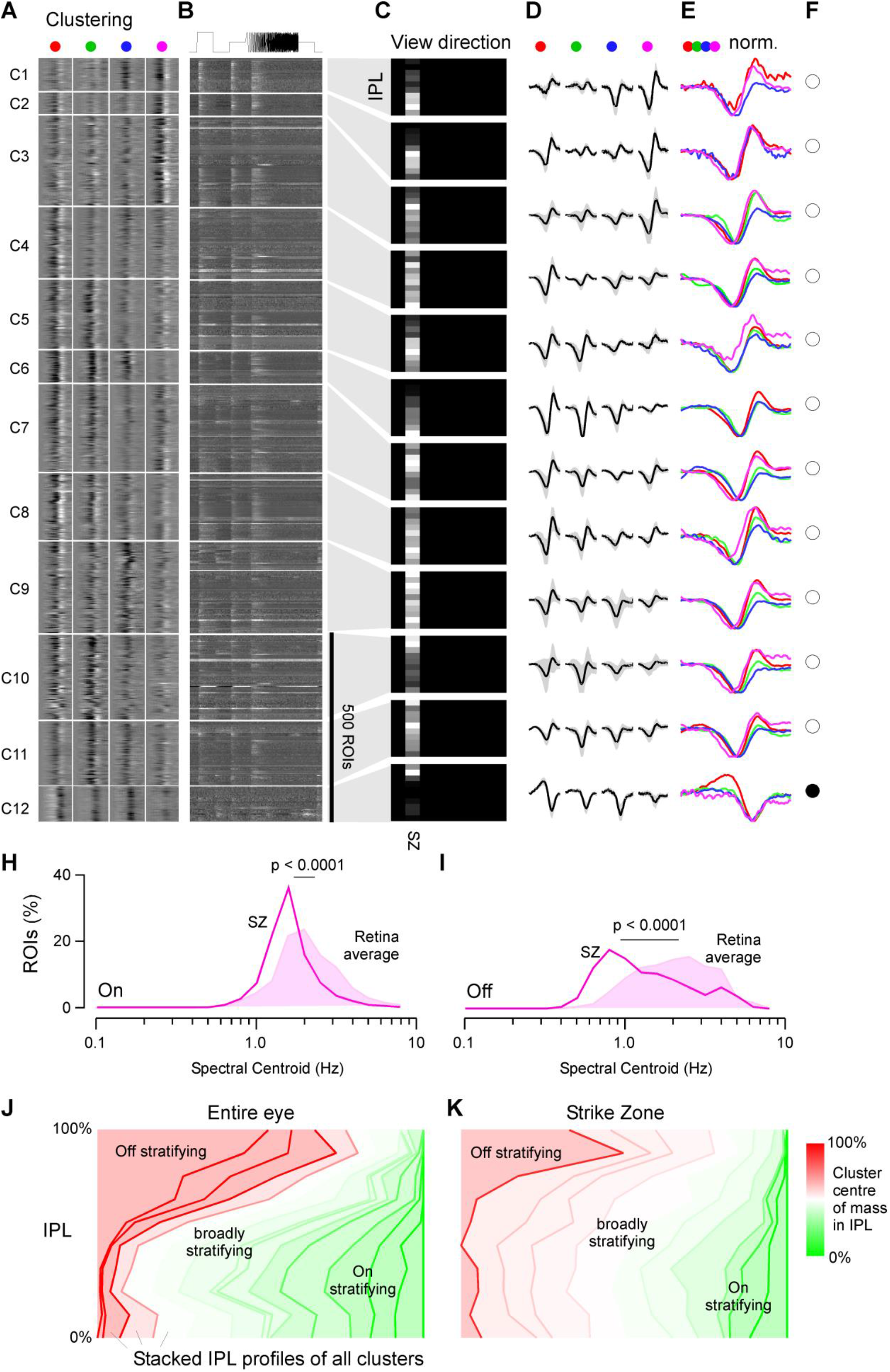
The SZ is dominated by broadly stratifying UV-sensitive On-clusters. **A-F**, Clustering of dendritic ROIs from SZ dataset (for details, cf. Fig. 6A-G). Note that all clusters except on (C_12_) are dominated by On-kernels, with C_1-3_ showing dominant UV-responses despite the relatively low UV-signal power in the stimulation light (cf. Fig. 2B). For corresponding clustering of SZ-somata, see Supplementary Fig. S4. **H,I**, Relatively slowed central frequency tuning of SZ-UV kernels (lines) compared to the retina average of UV-kernels (filled) amongst both On (H) and Off (I) kernels (cf. Fig. 4L,K). Both p<0.0001, Wilcoxon Rank Sum test, 1 tailed. **J,K**, Side-to-side comparison of functional stratification profiles of clusters from data across the eye (J, cf. Fig. 6C) and from SZ-only (K, cf. Fig. 8C). In each case, all cluster stratification profiles of a dataset were sorted by their centre of mass in the IPL (from 100%: Off to 0%: On), stacked on top of each other, and normalised to the number of ROIs per IPL depth. In addition, profiles were colour-coded by their centre of mass in the IPL as indicated. Note that most SZ-clusters (K) tended to broadly cover much of the IPL with a centre of mass near the middle of the IPL (white), while eye-wide stratification profiles (J) instead showed a greater tendency to stratify in either Off (red) or On (green) layers.

### RGC circuits in the strike zone

Following the same experimental approach as before (Fig. 2), we recorded from an additional 3,542 dendritic and 1,694 somatal ROIs in the SZ (Fig. 7A) of which 2,435 (69%) and 721 (42.6%), respectively, passed our response quality criterion (n=87 scans, 28 fish). In line with our whole-eye data (cf. Fig. 3–6), RGCs in the SZ were strongly On-biased across all wavelengths, including even a slight On-bias amongst blue responses (Fig. 7B,C, Fig. 8). In particular, SZ circuits exhibited a marked increase the abundance of UV-On responses, which were now also a dominant feature of several functional clusters (Fig. 7D,E; dendritic C_1-3_, Fig. 8, cf. Supplementary Fig. S4). Diverse RGC-functions mixed UV-On components with a variety of spectral and temporal non-UV components, which in most cases resulted in a spectrally biased but broad On-response (Fig. 7D,E, Fig. 8). Moreover, SZ UV-kernels were generally slower compared to the remainder of the eye (Fig. 8H) – in line with prolonged integration times of UV-cones in this part of eye for supporting prey capture (Yoshimatsu et al., 2019).

The diversity of both short- and long-wavelength biased On-circuits in addition to broadly tuned circuits might be helpful to support the detection of brighter-than-background prey objects in a variety of spectral lighting conditions, and might go partway to explaining why prey capture behaviour and associated brain activity can incur even in the absence of UV-illumination (Antinucci et al., 2019; Bianco et al., 2011; Mearns et al., 2019; Semmelhack et al., 2014) or indeed the absence of UV-cones (Yoshimatsu et al., 2019). Finally, a minority (~5%) of ROIs were allocated to a single, long-wavelength biased Off-cluster (C_12_). Such Off-circuits might underlie the detection of darker-than-background (prey-) objects (Bianco et al., 2011), which leads to the testable prediction that in this case UV-light should only play a minor role in behavioural performance.

SZ RGC-circuits did however not only functionally differ from those observed in the remainder of the eye. In addition, they also appeared to differ in their overall anatomical distribution across the depth of the IPL: SZ-RGC clusters appeared to be more broadly stratified (Fig. 8J,K). In fact, the only functional cluster that exhibited a narrow distribution across the IPL was the single OFF cluster C_12_ (Fig. 8K, cf. Fig. 8C). A broad stratification strategy amongst SZ ON-circuits might be useful to integrate retinal signals across a broad range of presynaptic circuits that encode a common position in visual space. Conceptually, such an arrangement might be a key requisite to build high signal-to-noise RGC circuits with small receptive fields for reliable detection of small targets during prey capture (Bianco et al., 2011; Semmelhack et al., 2014; Yoshimatsu et al., 2019).

Taken together, RGC circuits in the SZ exhibit therefore a key mix of response properties that may prove to be invaluable for visually guided prey capture of small and UV-bright paramecia (Yoshimatsu et al., 2019). Nevertheless, our findings thus far derive from population data of multiple RGCs that are superimposed in individual scan fields. To elucidate the identity of candidate RGC types underlying these specialisations, we therefore next turned to targeting individual RGCs (Figs. 9–11).

**Figure 9.**
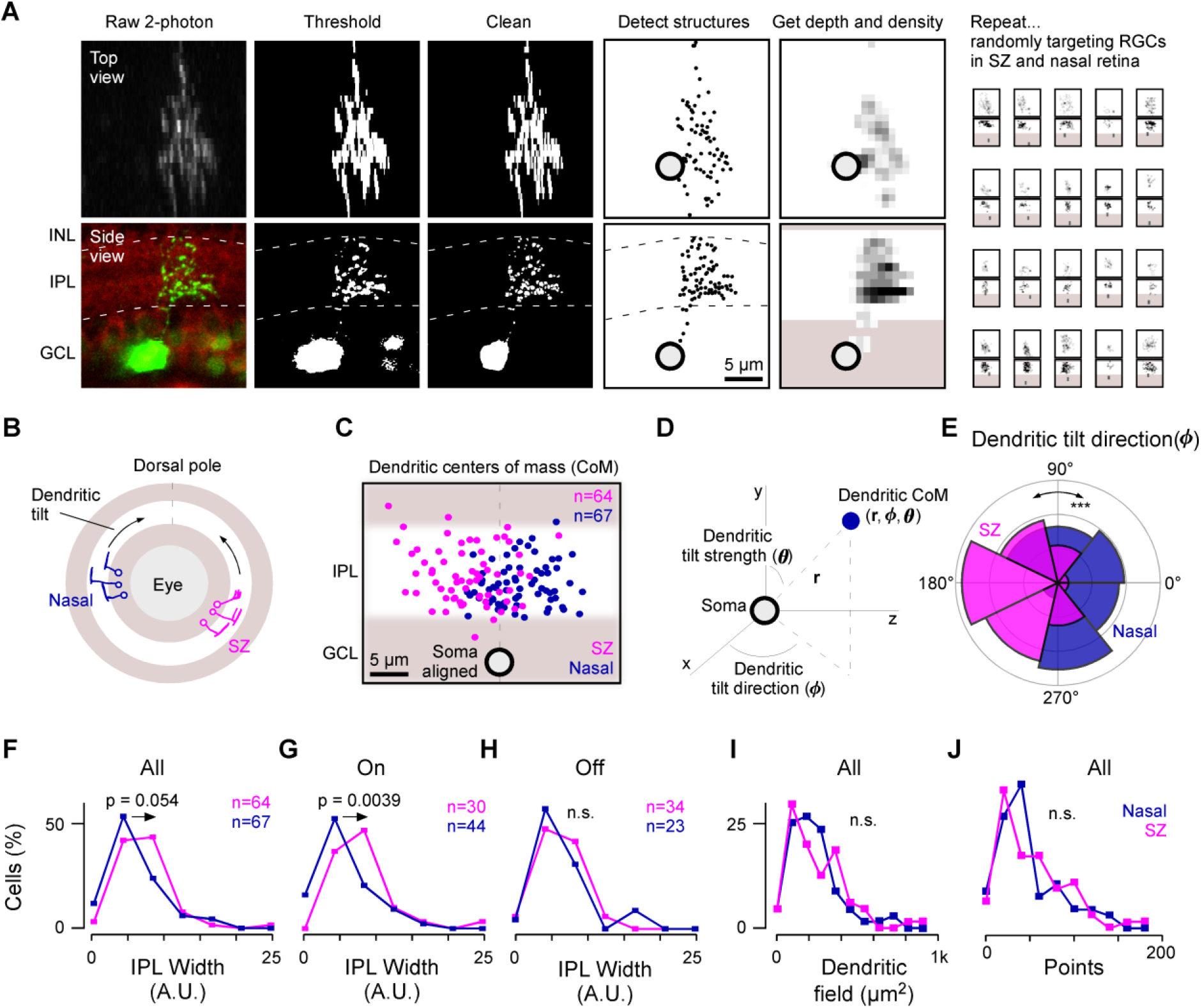
A relative overrepresentation of diffuse ON-RGCs in the SZ. **A**, Illustration of photoconversion and pre-processing pipeline for digitizing single RGC morphologies. Following photoconversion, cells were imaged as stacks under 2-photon (green) in the background of BODIPY staining to demarcate the IPL borders (red). Cells were thresholded and manually ‘cleaned’ where required prior to automatic detection of image structures and alignment relative to the IPL borders. Finally, the resultant ‘point clouds’ were used to determine summary statistics of each cell, and were also projected into density maps for visualisation. **B-J**, A total of n = 64 and n = 67 randomly targeted RGC from the SZ and nasal retina, respectively, were processed for further analysis, which included computation of their dendritic tilt (B-E), stratification widths within the IPL (F-H), en-face dendritic field area (I) and total number of detected dendritic structures (‘points’, J) (Methods). The dendrites of nasal (purple) and SZ (pink) RGCs both tended to tilt towards the eye’s dorsal pole (B: schematic, C: soma-aligned data of all dendrites’ centre of mass). Dendritic tilt was quantified in soma-centred polar coordinates based on the cartesian x,y,z coordinates that emerge from the original image stacks (D), such that r: distance in microns between soma and dendritic centre of mass (Supplementary Fig. S5A), θ (0:90°): strength of the dendritic tilt (0 and 90° denoting no tilt and maximal positive tilt, respectively, Supplementary Fig. S5B), and φ (0:360°): direction of the dendritic tilt in approximately retinotopic space (approximate as the eye is curved). φ Significantly differed between nasal and SZ RGCs (E). For summarising widths, RGC were considered as a single group (F) or split into On- and Off-RGCs (G, H, respectively), based on the IPL depths of their dendritic centre of mass (here the upper third of the IPL was considered “Off”, while the bottom two-thirds were considered “On”). Kolmogorov-Smirnov test for circular statistics (E) and Wilcoxon Rank Sum test, 1 tailed (F-J).

**Figure 10.**
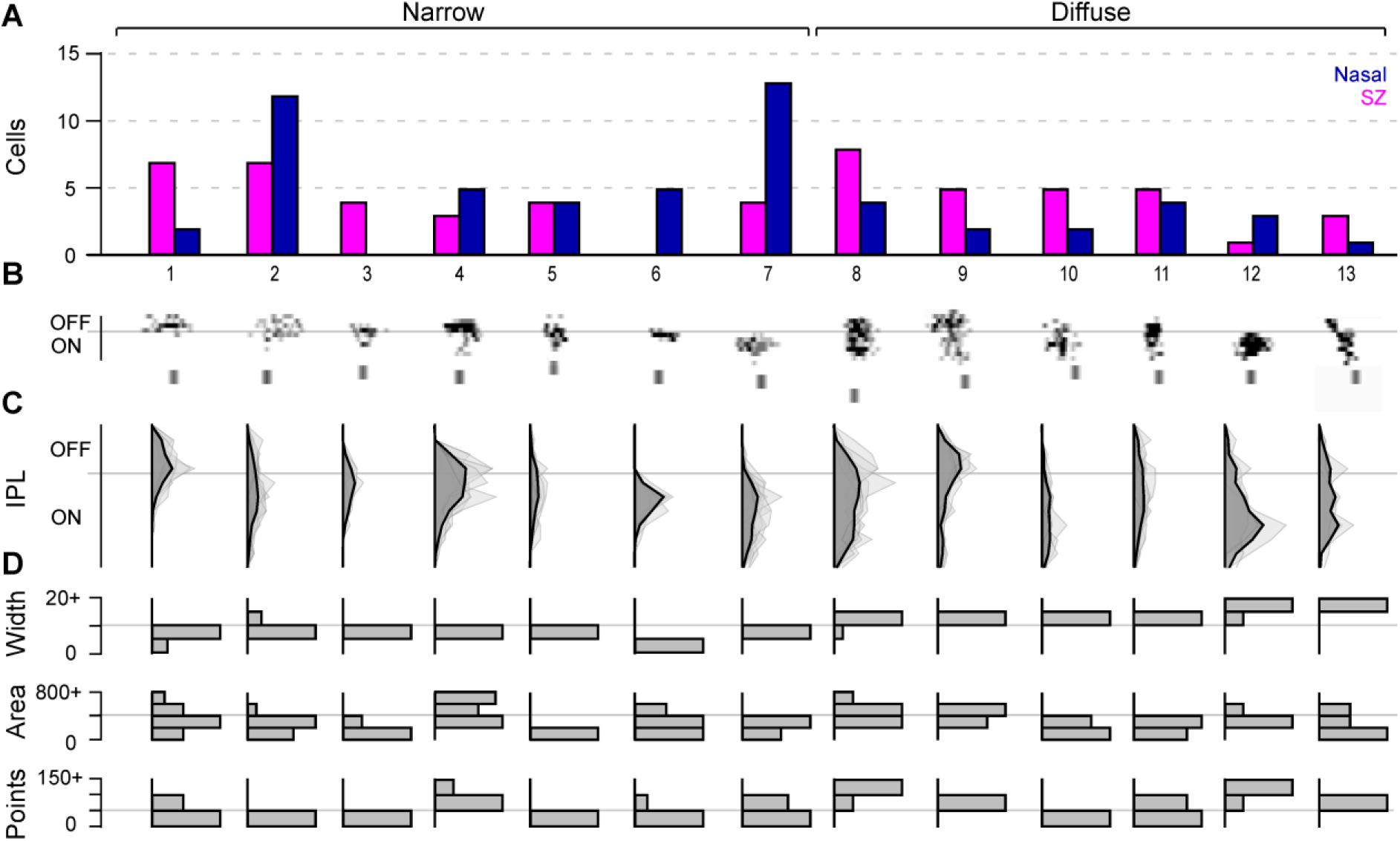
An asymmetric distribution of anatomical RGC types across the eye. Photoconverted and processed RGCs from both nasal and SZ (cf. Fig. 9) were jointly clustered based on morphological criteria (Methods). **A**, Number of RGCs for SZ (pink/ left) and nasal retina (blue/right) allocated to each of n = 13 clusters. **B**, Individual RGC morphologies representative for each cluster. Note that each morphology’s depth profile (y) is stretched five-fold relative to its lateral spread (x) to highlight stratification differences between clusters. **C**, Mean (dark) and individual depth profiles (light) and **D**, distribution of widths, dendritic field area and number of puncta for each cluster. Clusters were divided into narrow (left) and diffusely stratified (right) based on their mean widths (D, top, cf. labels in A).

**Figure 11.**
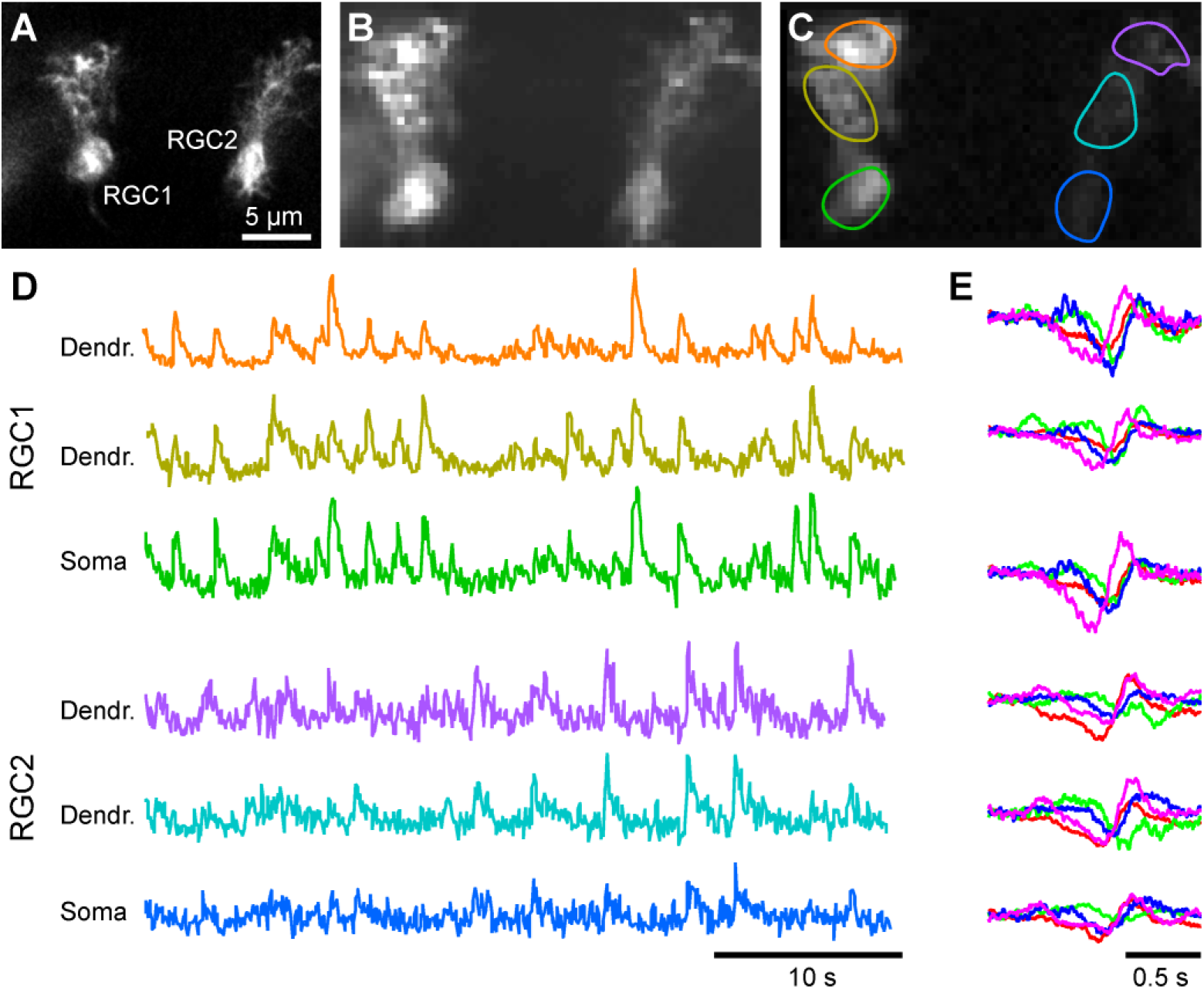
Putative ‘prey-capture-RGCs’ revealed following sparse expression. **A-C**, High-resolution scan of two randomly mGCAMP6f-expressing small-field but diffusely stratifying RGCs in the SZ that match our search terms for putative ‘prey-capture-RGCs’ in larval zebrafish (A), with activity scan (B) and correlation projection following visual stimulation with manually drawn ROIs (C). **D,E**, Example fluorescence traces from the ROIs in (C) in response to tetrachromatic white noise stimulation (D, stimulus not shown) and corresponding spectral kernels (E).

### Different morphological RGC types inhabit different parts of the eye

To assess the morphology of individual RGCs across the retina in an unbiased manner, we expressed photoactivatable (PA)-GFP (Patterson and Lippincott-Schwartz, 2002) in RGCs (Methods). Individual GCL somata were photoconverted (Fig. 9A, Methods) at random in two regions of the eye: SZ and nasal retina (N). A total of n = 222 RGCs from n = 113 fish were converted and imaged. After discarding n = 3 dAC which has had no obvious axon, and another n = 88 RGCs which were either incompletely labelled or which overlapped with neighbouring labelled RGCs, a final total n = 64 (SZ) and n = 67 (N) single RGCs were retained for further analysis. From here, we semi-automatically detected each RGC’s dendritic swellings as proxies for synaptic structures (Methods), and computed each swelling’s 3-dimensional location within the boundaries of the IPL, as determined after BODIPY counterstaining (Franke et al., 2017; Robles et al., 2014). We then used the resultant 3D ‘point-clouds’ to build depth and density profiles of each RGC for basic visualisation (Fig. 9A, right) and to extract basic metrics about their morphology: Specifically, for each RGC we computed the degree and direction of spatial offset between their soma and dendrites (‘dendritic tilt’, Figs. 9B-E), stratification width (narrow or diffuse, Fig. 9F-H), *en-face* dendritic area (Fig. 9I), and their total number of dendritic swellings (‘points’, Fig. 9J). Together, this revealed systematic morphological differences between RGCs randomly sampled from the SZ and nasal retina.

First, and in contrast to the majority of known RGC types in vertebrates (e.g.: (Bae et al., 2018; Dacey, 1999)) - the dendrites of most larval zebrafish RGC were spatially offset in retinotopic space relative to the position of their soma – reminiscent of ‘JamB’ (Joesch and Meister, 2016; Kim et al., 2008) or ‘MiniJ’-type RGCs (Rousso et al., 2016) in mice. This ‘dendritic tilt’ consistently pointed towards the dorsal pole of the eye, thus resulting in retinotopically-opposite tilts amongst nasal and SZ RGCs (Fig. 9B-E, Supplementary Fig. S5). How this systematic asymmetry in larval zebrafish RGCs is set-up developmentally, and if and how it contributes to their function will be important to assess in the future.

Second, as predicted from our functional census (Fig. 8), On-, but not Off-stratifying SZ-RGCs tended to be more diffusely stratified across IPL depth than nasal RGCs (Fig. 9F-H), in line with the pronounced upwards-shift of the functional On-Off boundary and resultant ‘anatomical compression’ of Off-circuits in the SZ (cf. Fig. 4 and (Zimmermann et al., 2018)). However, in our limited sample there was no significant difference in the distribution of RGCs’*en-face* dendritic area (Fig. 9I) or numbers of dendritic swellings (Fig. 9J) between the two retinal regions.

We next asked to what extent these overall stratification differences between SZ and nasal On-RGCs (Fig. 9G) could be linked to the presence of distinct morphological types in different parts of the eye (Fig. 10). For this, we jointly clustered both SZ and nasal RGCs, taking into account their mean IPL depths, widths and number of swellings (Methods). This yielded a total of 25 morphological clusters of which 13 with a minimum of n = 4 individual members were considered for further analysis (Fig. 10A). In line with a previous manually annotated census (Robles et al., 2014), RGC clusters exhibited diverse dendritic profiles including a variety of both narrowly (C_1-7_) and diffusely stratified profiles (C_8-13_, Fig. 10B). However, several clusters were mostly made up of RGCs coming from only one of the two retinal regions (Fig. 10A). For example, narrowly stratified clusters C_2,6,7_ were dominated by nasal RGCs, while diffusely stratified clusters C_8-10,13_ instead mainly comprised SZ-RGCs. In fact, the only narrowly stratified clusters that were dominated by SZ-RGCs were Off-stratifying C_1,3_, again in line with the anatomical compression of functional SZ-Off circuits. Next, though numerically small, of particular note were clusters C_3,5,10,13_ which had the smallest dendritic areas (Fig. 10D). A small dendritic field is generally associated with a correspondingly small spatial receptive field (Jacoby and Schwartz, 2017) which would be critical to detect small prey-like visual targets (Bianco et al., 2011; Semmelhack et al., 2014; Yoshimatsu et al., 2019). In agreement, and despite the lack of statistical significance when comparing dendritic field areas across the entire dataset (Fig. 9I), three of these four clusters were dominated by SZ-RGCs. These clusters were also reminiscent of candidate “prey-capture-RGC” morphologies previously identified based on their central projections to axonal arborization field 7 (AF7) which is associated with prey capture (Antinucci et al., 2019; Semmelhack et al., 2014).

### Outlook: Linking single cell structure and function

Ultimately, a clear understanding of the structures and functions of specific RGC circuits underlying visual guided prey-detection/capture will likely require a specific genetic handle on individual RGC-types, which is currently unavailable. Nevertheless, by combining knowledge on the behavioural specificity to prey-like stimuli across visual space (Bianco et al., 2011; Mearns et al., 2019; Semmelhack et al., 2014; Yoshimatsu et al., 2019), functional interrogation of RGC circuits in the eye (Figs. 2–8) as well as the identification of regionally biased RGC morphologies that conceptually match the expected requirements for such circuits (Figs. 9,10) we might make useful headway in guiding our understanding of how this tiny animal can use its RGCs to detect, recognize and ultimately capture its unicellular prey. Summarising what we now know, we ought to be searching for a small number of potentially small-field but diffusely stratifying RGCs in the SZ that show a robust sustained On-response to UV-light, as we as possibly an additional On-response to longer wavelength light. To make one step in this direction, we transiently expressed GCaMP6f in random RGCs (Methods), screened for fish with expression in the SZ, and characterised labelled cells’ morphologies and light responses. Using this strategy, we encountered two RGCs in a single animal that fully matched our search terms (Fig. 11): Both were small-field RGCs that diffusely stratified across the majority of the IPL (reminiscent of anatomical cluster C_13_, Fig. 10), and both exhibited clear On-responses to UV stimulation as well as to longer wavelength light (reminiscent of C_1_ in Fig. 6 and C_1-3_ in Fig. 8). Notably, both RGCs showed highly correlated responses to tetrachromatic white noise stimulation across their dendritic tree and soma (Fig. 11D), tentatively suggesting that they may be electrotonically compact. Understanding if and how RGCs such as these indeed contribute to visual-prey capture behaviour will be an important goal in the future.

## DISCUSSION

We have shown that the structure, organisation and function of larval zebrafish RGC circuits depends strongly on their position in the eye – presumably to meet visuo-ecological and behavioural demands in their natural visual world (Baden et al., 2020). For example, the localised presence of sustained UV-On RGCs in the SZ (Figs. 6–8) can be linked to their behavioural requirement to detect and localise small UV-bright prey in the upper frontal visual field (Bianco et al., 2011; Mearns et al., 2019; Semmelhack et al., 2014; Yoshimatsu et al., 2019). Similarly, the dominance of long-over short-wavelength responses in the lower visual field (Fig. 3D) is likely related to the predominance of long-wavelength light in the lower water column (Muaddi and Jamal, 1991; Nevala and Baden, 2019), as well as the zebrafish’s behavioural need to monitor the ground beneath them for systematic image shifts that drive a long-wavelength biased optomotor response (Orger and Baier, 2005; Wang et al., 2020).

In these aspects, our data from RGCs builds on our previous findings on the spectral responses of presynaptic BCs (Zimmermann et al., 2018). However, not all functions of BCs were simply inherited by the downstream RGCs. For example, the striking dominance of slow blue-Off circuits amongst RGCs (Figs. 4–6) was not predicted from BCs, which instead displayed an approximately balanced mix of blue-On and -Off circuits (Zimmermann et al., 2018). The near-complete absence of blue-On signals in zebrafish RGCs is also in stark contrast to the importance of multiple blue-On RGC circuits in mammals (Marshak and Mills, 2014; Mills et al., 2014) including in primates (Calkins et al., 1998; Dacey, 1996; Dacey and Lee, 1994). Next, while many of the dominant spectral opponencies observed in RGCs (Fig. 5) are already present at the level of BCs (Zimmermann et al., 2018), RGCs tended to more obviously mix time and wavelength information (Fig. 4, 6). Surprisingly, there was no clear increase in the diversity of RGC functions (Fig. 6) compared to BCs (Zimmermann et al., 2018) – in contrast to the approximately three-fold increase in neurons types from BCs to RGCs in mice (Baden et al., 2018). It is however possible, and arguably likely, that this functional census of zebrafish RGC diversity would disproportionately increase if spatial processing were considered (Franke et al., 2019), which was not a focus of the present study.

### Linking wavelength to visual and behavioural functions

In general, our data from zebrafish supports the long-standing view that achromatic image forming vision in animals is dominated by mid- and long-wavelength channels (Figs. 3B-D, 4D-J) (Baden and Osorio, 2019). Though never demonstrated physiologically in a tetrachromat’s retinal output, a close link between mid/long-wavelength vision and achromatic vision has been discussed for diverse species of both invertebrates and vertebrates including humans (Buchsbaum and Gottschalk, 1983; Jacobs and Rowe, 2004; Osorio and Vorobyev, 2008; Solomon and Lennie, 2007). It allows visual systems to capitalise on the typically abundant presence of mid- and long-wavelength photons in natural light (Muaddi and Jamal, 1991) to support high spatial and temporal acuity vision carried by the majority of retinal channels (Atick et al., 1992; Baden et al., 2020; Buchsbaum and Gottschalk, 1983; Lewis and Zhaoping, 2006; Maloney, 1986). Spectral information can then be sent in parallel by a typically lower number of retinal output channels to ‘colour in’ the grayscale scene in central circuits (Baden and Osorio, 2019; Dacey and Packer, 2003; Jacobs, 1993; Kelber et al., 2003; Neitz and Neitz, 2011). A segregation of achromatic mid/long-wavelength vision and circuits for colour vision is arguably taken to the extreme in the eyes of many arthropods including fruit flies: Six of each ommatidium’s eight photoreceptors (R1-6) express the same mid-wavelength opsin to support achromatic image forming vision, while the remaining two photoreceptors (R7,8) in parallel provide information about contrasts in wavelength (Heath et al., 2020; Schnaitmann et al., 2018). Similarly, like many visual neurons in insects (Chen et al., 2019; Heath et al., 2020; Schnaitmann et al., 2018; Yang et al., 2004), the finding that also in zebrafish most opponent RGCs encode simple rather than complex opponencies is in line with previous work (Baden and Osorio, 2019; Kamermans et al., 1991, 1998; Zimmermann et al., 2018) and links to the predominance of simple-over complex spectral contrasts in natural scenes (Buchsbaum and Gottschalk, 1983; Lewis and Zhaoping, 2006; Maloney, 1986; Nevala and Baden, 2019; Ruderman et al., 1998; Zimmermann et al., 2018).

And yet, in case of larval zebrafish, this parsimonious textbook view on the basic organisation of circuits for colour vison in animals remains at odds with several further observations:

1. It does not explain why nearly half of all output channels are colour opponent – it should be substantially fewer (Lewis and Zhaoping, 2006; Nevala and Baden, 2019).
2. It does not explain the striking mix of time- and spectral information throughout the eye.
3. It does not explain the near complete absence of blue-On circuits or the pervasive presence and general slowness of the blue-Off channel.
4. It does not explain the complex distribution of diverse UV-responses throughout the eye.

One explanation for this series of apparent mismatches with dominant theories in colour vision might relate to an implicit assumption that most if not all spectral processing and opponency should in some way link to image forming colour vision (Baden and Osorio, 2019). However, spectral information can be useful in additional ways. For example, zebrafish might simply use two separate and spectrally distinct achromatic systems: One long-wavelength biased achromatic system for traditional image formatting vision, and a second, short-wavelength biased achromatic system to detect image features that happen to be particularly detectable in this waveband: prey and predators. Water strongly scatters UV-light (Janssen, 1981) which submerges the cluttered visual background in a horizontally homogenous UV-haze. Objects in the foreground, such as nearby paramecia or predators then stand out as UV-bright or UV-dark objects, respectively (Cronin and Bok, 2016; Yoshimatsu et al., 2019). This scatter of UV-light also sets up a profound vertical brightness gradient, thus providing a reasonable explanation of why UV-circuits mainly survey the upper visual field.

Such a hypothetical dual-achromatic strategy would leave the blue channel ‘stuck in between’, encoding a mixture of red/green background and the UV-foreground. As such, blue circuits could possibly provide a useful subtraction signal to better delineate achromatic red/green-vision from achromatic UV-vision.

Following this line of thought, if the purpose of blue-Off circuits were not primarily to support image forming colour vision but instead to serve as a universal background signal, we might disregard it from our account of colour-opponency in zebrafish RGCs (Figs. 5A, 7D, highlighted in brown): In this case, two of the three most abundant colour-opponent groups amongst both dendrites and somata (RG_On_-B_Off_ and RGU_On_-B_Off_) would be classed as non-opponent On-responses (Fig. 5A). The total fraction of remaining colour opponent RGCs would then drop to 28% and 32% amongst dendrites and somata, respectively – still abundant but decidedly more in line with our understanding of how animal colour vision systems are organised (Baden and Osorio, 2019).

In addition, opponency against blue-light might also serve other non-image-forming functions. For example, the spectrum of natural daylight changes in a predictable manner over the course of the day, and can thus serve as a timing cue for circadian entrainment (Lazopulo et al., 2019; Mouland et al., 2019; Walmsley et al., 2015). Similarly, because the spectrum of downwelling light varies predictably with increasing water depth, spectral opponency can also serve as a depth gauge (Verasztó et al., 2018).

Finally, the striking link between spectral and temporal processing might also be reasonably explained by a dual-achromatic strategy segregated by a blue channel: A blue-Off background subtraction system might benefit from a long integration time to be relatively less perturbed by rapid changes in the visual scene. In contrast, UV-circuits for predator detection might then benefit from short integration times to trigger rapid escape behaviours, while UV-circuits for prey detection might benefit from slow-integration times to temporally accumulate evidence on the presence or absence of tiny prey items. Red/green systems might instead benefit from intermediate integration times to balance the need to faithfully encode both spatial and temporal aspects for traditional image forming vision.

Notwithstanding, these ideas - which conceptually may operate either in parallel to or intermixed with traditional circuits for image-forming colour vision - remain largely speculative. In the future it will be important to specifically explore testable predictions that emerge. For example, a selective presence or absence of blue light amongst otherwise natural spectrum white light illumination should markedly affect visual brain-circuit functions and behavioural performance during the presentation of conflicting visual stimuli that drive short- or long-wavelength dependent visual circuits (e.g. prey capture vs. optomotor reflex).

### The zebrafish area temporalis as an accessible model for the primate fovea?

Much of our own visual experience and central visual processing is dominated by the fovea, and regional damage to this part of the retina can be devastating to our sense of sight and quality of life (Bringmann et al., 2018). Accordingly, studies investigating the function and dysfunction of foveal circuits are of broad interest. However, most accessible model systems in vertebrate vision research, notably including mice, do not feature a similar specialisation (Baden et al., 2020). As a result, most studies on foveal function and dysfunction have remained restricted to primates, which dramatically limits options for experimental manipulations. However, the larval zebrafish’s *area temporalis* (strike zone, SZ) mimics several properties of the primate fovea, and may thus serve as a potentially useful and experimentally accessible alternative. Behaviourally, larval zebrafish specifically guide their SZ onto prey targets during fixational eye movements for high acuity binocular vision and distance estimation (Bianco et al., 2011; Mearns et al., 2020; Yoshimatsu et al., 2019), in many ways similar to fixational eye-movements in primates. Functionally, zebrafish SZ UV-cones boost signal-to-noise by using enlarged outer segments and slowed kinetics based molecular tuning of their phototransduction cascade (Yoshimatsu et al., 2019) - all specialisations that also occur in primate foveal cones (Peng et al., 2019; Sinha et al., 2017). Here, our data on RGC distributions and functions in larval zebrafish lends further credence to this notion. First, zebrafish have a fovea-like reduced AC to RGC ratio in their SZ (Fig. 1). Second, like in the primate fovea (Dacey, 2000; Sinha et al., 2017), SZ RGC circuits are spectrally distinct those of the peripheral retina (Fig. 4,6), and they are also slower (Fig. 8H, I). Third, retinal ganglion cells in the SZ are structurally distinct from those located in rest of the eye (Fig. 9) and include anatomical types that have a tiny dendritic field area that barely exceeds the width of their soma (Figs. 10,11) – the latter could conceivably form the substrate for a fovea-like extremely low-n convergence of BC and cone signals. In the future it will be interesting to explore what further aspects of the zebrafish SZ – if any – can be paralleled to foveal vision in primates. Moreover, it will be critical to evaluate to what extent this growing series of functional, structural and molecular links between the two retinal systems may generalise across acute zones of other vertebrates (Baden et al., 2020).

## METHODS

### Data availability

Pre-processed functional data as well as single-RGC morphological data, associated summary statistics, cluster allocations (where applicable) and basic analysis and clustering scripts written in Matlab and can be accessed from DataDryad via the relevant links on www.retinal-functomics.net. Any remaining data will be provided upon reasonable request to the corresponding author.

### Animals

All procedures were performed in accordance with the UK Animals (Scientific Procedures) act 1968 and approved by the animal welfare committee of the University of Sussex. Adult animals were housed under a standard 14/10 light/dark cycle and fed 3 times daily. Larvae were grown in E2 solution (1.5M NaCl, 50mM KCl, 100mM MgSO_4_, 15mM KH_2_PO_4_, 5mM Na_2_HPO_4_) or fish water and treated with 200μM 1-phenyl-2-thiourea (PTU: Sigma) from 12 hours post fertilization (*hpf*) to prevent melanogenesis (Karlsson et al., 2001). For 2-photon in-vivo imaging, zebrafish larvae were immobilised in 2% low melting point agarose (Fisher Scientific, BP1360-100), placed on a glass coverslip and submerged in fish water. Eye movements were prevented by injection of α-bungarotoxin (1 nL of 2 mg/ml; Tocris, Cat: 2133) into the ocular muscles behind the eye.

For all experiments, we used 6-8 *dpf* zebrafish (*Danio rerio*) larvae. The following previously published transgenic lines were used: *Tg(Ptf1a:dsRed)* (Jusuf and Harris, 2009), *Tg(Islet2b:nls-trpR, tUAS:MGCamp6f)* (Janiak et al., 2019) as well as Casper (White et al., 2008), nacre (Thisse et al., 1993) and roy (Ren et al., 2002). In addition, two transgenic lines *Tg(Islet2b:nls-trpR, tUAS:SyjRGeco1a)*and *Tg(tUAS:paGFP)* were generated by injecting plasmid solution into one-cell stage embryos. Plasmid solution used are; a mixture of pTol2pA-islet2b-nlsTrpR (Janiak et al., 2019) and pTol2CG2-tUAS-SyjRGeco1a for the *Tg(islet2b:nls-trpR, tUAS:SyjRGeco1a)* line and pTol2BH-tUAS-paGFP for the *Tg(tUAS:paGFP)*line. Expression of paGFP was then obtained by crossing these two lines. With this combination, RGCs also express SyjRGeco1a, which was not used in this study (and which did not interfere with the green channel used for paGFP detection.

Plasmids were constructed by means of a attL/attR (LR)-reaction using destination and entry plasmids as follows; for pTol2CG2-tUAS-SyjRGeco1a; pDestTol2CG2 (REF), p5E-tUAS(REF), pME-SyjRGeco1a, p3E-pA(REF), for pTol2BH-tUAS-paGFP; pDestTol2BH (REF), p5E-tUAS, pME-paGFP, p3E-pA. pME-SyjRGeco1a was constructed by inserting PCR amplified zebrafish synaptophysin without stop codon (Dreosti et al., 2009) followed by PCR amplified jRGeco1a fragment (Dana et al., 2016) into pME plasmid. Similarly, pME-paGFP was constructed by inserting PCR amplified paGFP fragment into pME plasmid.

For transient expression of mGCaMP6f under Islet2b we injected a mixture of pTol2pA-islet2b-nlsTrpR and pTol2BH-tUAS-MGCamp6f plasmids solution into one-cell stage eggs. Positive embryos were screened under 2-photon.

### Tissue preparation, immunolabeling, and imaging

For immunohistochemistry, larvae were euthanised by tricaine overdose (800 mg/l) and fixed in 4% paraformaldehyde in phosphate-buffered saline (PBS) for 30 minutes at room temperature before being washed in calcium-negative PBS. Retinae were then incubated in permeabilization/blocking buffer (PBS with 0.5% Triton X-100 and 5% normal donkey serum) at 4°C for 24 hours, and thereafter transferred to the appropriate labelling solution. For nuclear labelling, tissue was incubated at 4°C in blocking solution with Hoechst 33342 nuclear dye (Sigma, H21492, 1:2000) for 24 hours. For membrane staining, tissue was incubated at 4°C in blocking solution with BODIPY membrane dye (Sigma, D3821, 1:1000) for 24 hours. For immunostaining, tissue was incubated at 4°C for 72 hours in primary antibody solution (chicken anti-GFP (AbCam, 13970, 1:500), rabbit anti-cox iv (AbCam, 16056, 1:500), diluted in permeabilization/blocking solution). Samples were rinsed three times in PBS with 0.5% Triton X-100, then transferred to secondary antibody solution (donkey anti-chicken IgG CF488 A conjugate (Sigma, 1:500), donkey anti-rabbit IgG CF568 conjugate (Sigma, 1:500)), diluted in permeabilization/blocking solution and incubated at 4°C for 24 hours. Finally, samples were rinsed three times in PBS with 0.5% Triton X-100 before being mounted in mounting media (VectaShield, Vector, H-1000) for confocal imaging.

GABA immunostaining was performed according to the protocol described by (Jusuf and Harris, 2009). Briefly, whole retinas were fixed in 2% PFA /2%glutaraldehyde for 24 hours at 4°C, rinsed in PBS, treated with 0.1% sodium borohydride (NaBH_4_) in 0.2% Triton X-100 in PBS for 10 minutes at room temperature, and rinsed again to remove excess NaBH4. For immunolabeling, all steps are as described above, with the following exceptions: blocking buffer consisted of 10% normal donkey serum, 0.1% Tween-20, and 0.5% Triton X-100 in PBS; primary and secondary antibodies were also diluted in this blocking buffer.

Confocal stacks and individual images were taken on Leica TCS SP8 using 40x water-immersion objective at xy resolution of 2,048×2,048 pixels (pixel width: 0.162 μm). Voxel depth of stacks was taken at z-step 0.3-0.5 μm. Contrast and brightness were adjusted in Fiji (NIH).

### Cell density mapping

The 3D positions of all GCL somata (stained with Hoecht 3342), as well as dAC and AC somata *(tg(Ptf1a:dsRed)*, and MG *tg(GFAP:GFP)*, immunolabeled against GFP) were semi-automatically detected in Fiji from confocal image stacks of intact, whole eyes. These positions were then projected into a local-distance preserving 2D map as shown previously (Zimmermann et al., 2018) using custom-written scripts in Igor Pro 6.37 (Wavemetrics). The density map of RGC somata was computed by subtracting the density map of dACs from that of GCL cells. Similarly, the density map of ACs was computed by summing the density maps of dACs and ACs from the inner nuclear layer. From here, RGC maps were also mapped into a sinusoidal projection of visual space (Yoshimatsu et al., 2019).

### Axonal tracing

The lipophilic tracer dye DiO (Invitrogen, D307) was used to trace RGC axons from the retina to their arborization fields in the pretectum and tectum. 1 mg/mL stock solution was prepared in dimethylformamide and stored at −20°C. For injection into *Tg(Islet2b:nls-trpR, tUAS:MGCamp6f)* retinas, the lenses of whole fixed larvae were removed and a sufficient amount of tracer dye injected into one of either the left or the right eye so as to completely cover the exposed surface of the GCL. Tissue was then incubated at 37°C for 3 days to allow the dye time to diffuse all the way up RGC axons to their terminals in the midbrain.

### Photoactivation

Prior to photoconversion, 6-8 *dpf* Islet2b:PA-GFP larvae were injected with BODIPY membrane dye (1nL of 1mg/mL; Sigma, D3821) into the space behind the right eye and underlying skin to demarcate retinal anatomy and facilitate subsequent targeting. Larvae were left for 10-20 minutes at 25°C to allow the dye to diffuse into the retina. After 20 minutes, the IPL was uniformly stained, and the individual somata of GCL neurons showed nuclear exclusions which were used for subsequent targeting.

Cells were photoconverted under the same 2-photon microscope as used for functional imaging (below). In each animal, we randomly photoconverted 2-5 cells per eye in the nasal retina and/or strike zone, with a minimum spacing of 30 μm between them. For photoactivation, the femtosecond laser was tuned to 760 nm and focused onto one single soma at a time for up to ~2 minutes. After a typically >40 minutes cells were visualised under 2-photon (927 nm) and imaged in a 512×512 pixel (1 μm z-steps) stack which encompassed each cell’s soma, axon initial segment, and the entirety of the dendritic structure. Throughout, the BODIPY signal was included as an anatomical reference.

### Two-photon functional imaging and stimulation parameters

For all *in vivo* imaging experiments, we used a MOM-type two-photon microscope (designed by W. Denk, MPI, Martinsried (Euler et al., 2013); purchased through Sutter Instruments/Science Projects) equipped with the following: a mode-locked Ti:Sapphire laser (Chameleon Vision-S, Coherent) tuned to 927 nm for imaging GFP and 960 nm for imaging mCherry/BODIPY in combination with GFP; two fluorescent detection channels for GFP (F48×573, AHF/Chroma) and mCherry/BODIPY (F39×628, AHF/Chroma), and; a water-immersion objective (W Plan-Apochromat 20x/1,0 DIC M27, Zeiss). For image acquisition, we used custom-written software (ScanM, by M. Mueller, MPI, Martinsried and T Euler, CIN, Tübingen) running under Igor Pro 6.37 (Wavemetrics). Structural data was recorded at 512×512 pixels, while functional data was recorded at 64×32 pixel resolution (15.6 Hz, 2 ms line speed). For each functional scan, we first defined a curvature of the imaged IPL segment based on a structural scan, and thereafter “bent” the scan plane accordingly (“banana scan”). This ensured that the imaging laser spent a majority of time sampling from the curved IPL and INL, rather than adjacent dead-space. The banana-scan function was custom-written under ScanM.

For light stimulation, we focused a custom-built stimulator through the objective, fitted with band-pass-filtered light-emitting diodes (LEDs) (‘red’ 588nm, B5B-434-TY, 13.5 cd, 8°; ‘green’ 477 nm, RLS-5B475-S, 3-4cd, 15°, 20 mA; ‘blue’ 415 nm, VL415-5-15, 10-16mW, 15°, 20 mA; ‘ultraviolet’ 365 nm, LED365-06Z, 5.5 mW, 4°, 20 mA; Roithner, Germany). LEDs were filtered and combined using FF01-370/36, T450/pxr, ET420/40 m, T400LP, ET480/40x, H560LPXR (AHF/Chroma). The final spectra approximated the peak spectral sensitivity of zebrafish R-, G-, B-, and UV-opsins, respectively, while avoiding the microscope’s two detection bands for GFP and mCherry/BODIPY. To prevent interference of the stimulation light with the optical recording, LEDs were synchronized with the scan retrace at 500Hz (2 ms line duration) using a microcontroller and custom scripts. Further information on the stimulator, including all files and detailed build instructions can be found at https://github.com/BadenLab/Tetra-Chromatic-Stimulator.

Stimulator intensity was calibrated (in photons per second per cone) such that each LED would stimulate its respective zebrafish cone type with a number of photons adjusted to follow the relative power distribution of the four wavelength peaks of daytime light in the zebrafish natural habitat (Nevala and Baden, 2019; Zimmermann et al., 2018) to yield ‘natural white’: red, “100%” (34×10^5^ photons /s /cone); green, “50%” (18 x 10^5^ photons /s /cone); blue, “13%” (4.7 x 10^5^ photons /s /cone); ultraviolet, “6%” (2.1×10^5^ photons /s /cone). We did not compensate for cross-activation of other cones. Owing to 2-photon excitation of photopigments, an additional constant background illumination of ~10^4^ R* was present throughout (Baden et al., 2013; Euler et al., 2009, 2019). For all experiments, larvae were kept at constant illumination for at least 2 seconds after the laser scanning started before light stimuli were presented. Two types of full-field stimuli were used: a binary dense “natural spectrum” white noise, in which the four LEDs were flickered independently in a known random binary sequence at 6.4Hz for 258 seconds, and a natural-white chirp stimulus (Baden et al., 2016) where all four LEDs were driven together. To prevent interference of the stimulation light with the optical recording, LEDs were synchronised to the scanner’s retrace (Euler et al., 2019).

### Quantification and statistical analysis

No statistical methods were used to predetermine sample size.

### Data analysis

Data analysis was performed using IGOR Pro 6.3 (Wavemetrics), Fiji (NIH) and Matlab R2018b (Mathworks).

### ROI placements and quality criterion

ROIs were automatically placed using local image correlation based on established protocols – for details see (Franke et al., 2017). To allocate ROIs to dendritic and somatic datasets a boundary between the GCL and IPL was drawn by hand in each scan - all ROIs with a centre of mass above the boundary were considered as dendritic, and all ROIs below were considered as somatic. Due to the ring-like nature of mGCaMP6f expression profiles in somata when optically sectioned, it was possible that two ROIs could be inadvertently placed on different halves of the same soma. Since whether or not a soma was split in this way was likely non-systematic over functional types, we did not attempt to correct for this possibility. Only ROIs where at least one of the four spectral kernels’ peak-to-peak amplitudes exceeded a minimum of ten standard deviations were kept for further analysis (n = 2,716/2,851 dendritic ROIs, 95%; 586/796 somatal ROIs, 74%). Equally, all individual colour kernels that did not exceed 10 SDs were discarded (i.e set to NaN).

### Kernel polarity

The use of a fluorescence-response-triggered average stimulus (here: ‘kernel’) as a shorthand for a neuron’s stimulus-response properties, while potentially powerful (e.g. (Franke et al., 2017; Szatko et al., 2019; Zimmermann et al., 2018)), ought to be considered with some caution. For example, determining a binary value for a kernel’s polarity (On or Off) can be conflicted with the fact that a neuron might exhibit both On and Off response aspects. Moreover, different possible measures of On or Off dominance in a kernel can generate different classification biases. Here, we defined On and Off based on a measure of a kernel’s dominant trajectory in time. For this, we determined the position in time of each kernel’s maximum and minimum. If the maximum preceded the minimum, the kernel was classified as Off, while vice versa if the minimum preceded the maximum, the kernel was defined as On. Examples On and Off kernels classified by this metric can for example be seen in Fig. 5B (cf. Fig. 5A central horizontal column for a lookup of how each kernel was classified).

### Digitizing photoactivated cells

Dendritic swellings (taken as a proxy for synaptic densities) in photoconverted GCL cells were detected using Fiji. For this, the GFP channel was smoothed and thresholded to create a binary mask removing background fluorescence. Any remaining neurites that clearly did not belong to the most strongly labelled cell were removed by hand. Next, the soma and any dendritic swellings were automatically detected using 3D Objects Counter plugin in Fiji. 3D positions of all detected objects were then normalised relative to the boundaries of the IPL, as determined from the BODIPY channel. This generated an IPL-aligned 3D ‘dot-cloud’ for each RGC, which was then used as the input for a custom clustering algorithm. We also projected each dot-cloud into *en-face* and side-view density maps for visualisation. Note that sideview projections shown in Fig. 9A (rightmost) and Fig. 10B are laterally compressed five-fold to highlight differences in stratification depths across the IPL.

### Quantifying dendritic tilt

As noted in ‘Morphology Clustering’ (below), morphological data consists of sets of points in three-dimensional Cartesian coordinates (*x,y,z*) describing the location of the soma and the dendritic architecture for each RGC. The coordinate axes are orientated such that the *y*-axis is perpendicular to the plane of the retina, pointing outwards, away from the centre of the eye, while the *x* and *z* axes are tangential to the plane of the retina. We translated the coordinate system for each cell such that its soma lies at the origin. We then calculated the centre of mass (CoM) of the point cloud representing the dendritic tree of each cell (i.e. excluding the soma), computed as the mean of the points’ *x*, *y* and *z* positions. We then transformed to a spherical polar coordinate system, (*r*, *θ*, *φ*), with the origin centred at the soma, where *r* > 0(μm) is the distance of the dendritic CoM from the soma, the polar angle 0 ≤ *θ* ≤ *π* (rad), characterises the dendritic tilt strength (i.e. the angle subtended by the dendritic CoM from the *y*-axis, where *θ* = 0 corresponds to no tilt and *θ* = *π*/2 occurs when the dendritic CoM has the same IPL/GCL depth as the soma) and the azimuthal angle, 0 ≤ *φ* < 2*π* (rad), characterises the dendritic tilt direction. It should be noted that the relationship between our Cartesian and spherical polar coordinate systems is different from that which is standard in that we have swapped the *y* and *z* axes. Thus, the polar angle is subtended from the *y*-axis, rather than from the *z*-axis as is usual.

We tested whether the distributions of the position of the dendritic CoM relative to the soma in each of the *r*, *θ* and *φ* dimensions for SZ and nasal RGCs are from the same (continuous) distribution using the two-sample Kolmogorov-Smirnov test. This was implemented using the Matlab routine **kstest2** for *r* and *θ*, and using the **circ_kuipertest** routine from the CircStat toolbox (Berens, 2009) for *φ*, since this variable is (2*π*-) periodic. In comparing SZ and nasal RGCs, the dendritic CoM positions, *r*, are predicted to be from different distributions (*p* = 0.0209, 3 s.f.); the dendritic tilt strengths, *θ*, are predicted to be from the same distribution (*p* = 0.894, 3 s.f.); and the dendritic tilt angles, *φ*, are predicted to be from different distributions (*p* = 0.001).

### Morphology Clustering

The morphological data consists of sets of points in three-dimensional Cartesian coordinates (x,y,z) describing the dendritic architecture for each of 131 RGCs, 67 from the nasal (N) region and 64 from the strike zone (SZ) region. The coordinate axes are orientated such that the y-axis is perpendicular to the plane of the retina, spanning the width of the IPL, while the x and z axes are tangential to the plane of the retina. The coordinates in the y-dimension are scaled so as to lie in the interval [0,10] for any processes within the IPL, and beyond >10 or <0 or for INL and GCL processes (where applicable), respectively. The position of the soma, which always lay in the GCL, was not used for clustering.

Three summary statistics, each of which capture some aspect of the dendritic architecture, were defined for use in clustering: i) y_span: the width of the dendritic tree in the y-direction; ii) y_mean: the mean position of the points in the dendritic tree in the y-direction; and iii) num_pts: the number of points in the dendritic tree. While we experimented with other summary statistics, these three were found to be sufficient to differentiate the RGCs into their basic morphological groups.

We also defined one further summary statistic: iv) xz_area: the area spanned by the dendritic tree in the xz-plane, calculated as the convex hull using the Matlab routine **convhull**. This statistic was not used for clustering since the information contained in xz_area is largely captured between y_span and num_pts. While not required for clustering, this summary statistics nonetheless captures important characteristics of the dendritic morphology and hence is represented in the results section alongside y_span, y_mean and num_pts.

Each of the summary statistics was standardised by subtracting the mean and dividing by the standard deviation. In this way, we ensured that each of the summary statistics was equally weighted by the clustering algorithm.

Clustering was performed in two stages, using agglomerative hierarchical clustering in both cases. The first stage of clustering used all three summary statistics (y_span, y_mean and num_pts), splitting the data into 18 clusters. Two of the resulting clusters were large and contained a variety of morphologies as discerned from visual inspection. These clusters were split further via a second round of clustering, using just the y_span summary statistic. The first cluster was split into 6 subclusters and the second into 3 subclusters, resulting in a total of 25 clusters, where the 13 clusters containing a minimum of 4 members were included for presentation.

Hierarchical clustering was performed using the Matlab routines **pdist**, **linkage** and **cluster**. The function **pdist** calculates the distances between each RGC in (y_span,y_mean,num_pts)-space, while the function **linkage** operates on the output of the **pdist** routine to encode an agglomerative hierarchical cluster tree. There are a number of options for defining the distances between RGCs for **pdist** and the distances between clusters for **linkage**. We used the ‘city block’ distance metric for **pdist** and the ‘average’ distance metric for **linkage** as, in general, these were found to result in a larger cophenetic correlation coefficient (CCC) than any other combination of distance metrics. The CCC is a measure of the fidelity with which the cluster tree represents the dissimilarities between observations. It was calculated using the Matlab routine **cophenet** and takes values between [−1,1], where values closer to positive unity represent a more faithful clustering. In the results presented here, the first stage of clustering had a CCC of 0.77 (2 d.p.), while the two subclusterings in the second stage had CCCs of 0.77 (2 d.p.) and 0.83 (2 d.p.).

Lastly, RGCs were assigned to clusters using the Matlab routine **cluster**. The number of clusters was determined by specifying a cutoff distance which was chosen following visual inspection of the cluster tree dendrogram so as to respect a natural division in the data.

### Functional data pre-processing and receptive field mapping

Regions of interest (ROIs), corresponding to dendritic or somatic segments of RGCs were defined automatically as shown previously based on local image correlation over time (Franke et al., 2017). Next, the Ca^2+^ traces for each ROI were extracted and detrended by high-pass filtering above ~0.1 Hz and followed by z-normalisation based on the time interval 1-6 seconds at the beginning of recordings using custom-written routines under IGOR Pro. A stimulus time marker embedded in the recording data served to align the Ca^2+^ traces relative to the visual stimulus with a temporal precision of 1 ms. Responses to the chirp stimulus were up-sampled to 1 KHz and averaged over 3-6 trials. For data from tetrachromatic noise stimulation we mapped linear receptive fields of each ROI by computing the Ca^2+^ transient-triggered-average. To this end, we resampled the time-derivative of each trace to match the stimulus-alignment rate of 500 Hz and used thresholding above 0.7 standard deviations relative to the baseline noise to the times *t_i_* at which Calcium transients occurred. We then computed the Ca^2+^transient-triggered average stimulus, weighting each sample by the steepness of the transient:

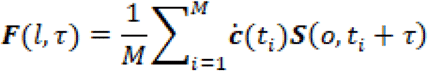

Here, *S*(*l, t*) is the stimulus (“LED” and “time”), *τ* is the time lag (ranging from approx. −1,000 to 350 ms) and *M* is the number of Ca^2+^ events. RFs are shown in z-scores for each LED, normalised to the first 50 ms of the time-lag. To select ROIs with a non-random temporal kernel, we used all ROIs with a standard deviation of at least ten in at least one of the four spectral kernels. The precise choice of this quality criterion does not have a major effect on the results.

### Eye-IPL maps

To summarise average functions of RGC processes across different positions in the eye and across IPL depths, we computed two-dimensional “Eye-IPL” maps. For this, we divided position in the eye (-π:π radians) into eight equal bins of width π/4. Similarly, we divided the IPL into 20 bins. All soma ROIs were allocated to bin 1 independent of their depth in the GCL. while all IPL ROIs were distributed to bins 3:20 based on their relative position between the IPL boundaries. As such, bin 2 is always empty, and serves as a visual barrier between IPL and GCL. From here, the responses of ROIs within each bin were averaged. All maps were in addition smoothed using a circular π/3 binomial (Gaussian) filter along eye-position, as well as for 5% of IPL depth across the y-dimension (dendritic bins 3:20 only).

### On-Off index (OOi)

For each Eye-IPL bin, an On-Off index (OOi) was computed:

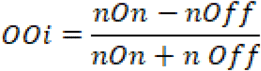

Where nOn and nOff correspond to the number of On and Off kernels in a bin, respectively. OOi ranged from 1 (all kernels On) to −1 (all kernels Off), with and OOi of zero denoting a bin where the number of On and Off kernels was equal.

### Ternary response classification

Each ROI was allocated to one of 81 ternary response bins (three response states raised to the power of four spectral bands). One of three response-states was determined for each of four spectral kernels (red, green, blue, UV) belonging to the same ROI: On, Off or non-responding. All kernels with a peak-to-peak amplitude below ten standard deviations were considered non-responding, while the remainder was classified as either On or Off based on the sign of the largest transition in the kernel (upwards: On, downwards: Off).

### Feature extraction and Clustering

Clustering was performed on four data sets, each containing the functional responses of RGCs to chirp stimuli and kernels derived from colour noise stimuli: 1) pan retinal inner plexiform layer (PR-IPL) data set (n = 2,851), sampling RGC responses at all eccentricities and across a range of depths in the IPL; 2) strike zone inner plexiform layer (SZ-IPL) data set (n = 3,542), sampling RGCs at the SZ only and across the IPL; 3) pan retinal ganglion cell layer (PR-GCL) data set (n = 796), sampling RGC responses at all eccentricities from the RGC somata in the GCL; and 4) strike zone ganglion cell layer (SZ-GCL) data set (n = 1,694), sampling RGCs at the SZ only from the RGC somata. Mean responses to chirp stimuli were formatted as 2,499 time points (dt = 1 ms) while colour kernels were formatted as 649 time points (dt = 2 ms, starting at t = −0.9735 s) per spectral channel (red, green, blue and UV).

For each dataset we clustered using only the kernels portion of the data since this was found to produce a cleaner clustering than when clustering chirp responses and kernels together, or chirp responses alone. ROIs with low quality kernels, determined as the maximum standard deviation across the four colours, were identified and removed from the data set. For clustering, a kernel quality threshold of 5 was chosen, such that any ROI with a kernel quality below this threshold was eliminated from the data to be clustered.

Following quality control, the data sets had the following sizes: 1) PR-IPL: n = 2,414 (84.7% of original); 2) SZ-IPL: n = 2,435 (68.8% of original); 3) PR-GCL: n = 411 (51.6% of original); 4) SZ-GCL: n = 721 (42.6 % of original).

We scaled the data corresponding to each kernel colour by dividing each one by the standard deviation through time and across ROIs. In this way we ensured an even weighting for each colour. This is important, since the red and green kernels tended to have larger amplitudes than the blue and UV kernels.

We used principal component analysis (PCA) to reduce the dimensions of the problem prior to clustering. PCA was performed using the Matlab routine **pca** (default settings). We applied PCA to the portions of a data set corresponding to each of the kernel colours separately, retaining the minimum number of principal components necessary to explain ≥99% of the variance. The resulting four ‘scores’ matrices were then concatenated into a single matrix ready for clustering. The following numbers of principal components were used for each of the four data sets: 1) PR-IPL: 8 red (R) components, 8 green (G) components, 13 blue (B) components, 33 ultraviolet (UV) components (62 in total); 2) SZ-IPL: 15 R, 17 G, 25 B, 18 UV (75 in total); 3) PR-GCL: 13 R, 11 G, 24 B, 36 UV (84 in total); and 4) SZ-GCL: 20 R, 21 G, 27 B, 34 UV (102 in total).

We clustered the combined ‘scores’ matrix using Gaussian Mixture Model (GMM) clustering, performed using the Matlab routine **fitgmdist**. We clustered the data into clusters of sizes 1,2,…,100, using i) shared-diagonal, ii) unshared-diagonal, iii) shared-full and iv) unshared-full covariance matrices, such that (100*4 =) 400 different clustering options were explored in total. For each clustering option 20 replicates were calculated (each with a different set of initial values) and the replicate with the largest loglikelihood chosen. A regularisation value of 10^-5^ was chosen to ensure that the estimated covariance matrices were positive definite, while the maximum number of iterations was set at 10^4^. All other **fitgmdist** settings were set to their default values.

In data sets PR-IPL and SZ-IPL the optimum clustering was judged to be that which minimised the Bayesian information criterion (BIC), which balances the explanatory power of the model (loglikelihood) with model complexity (number of parameters), while clusters with <10 members were removed. In data sets PR-GCL and SZ-GCL the BIC did not give a clean clustering; therefore, we specified 20 clusters for the PR-GCL and 10 clusters for the SZ-GCL, with unshared-diagonal covariance matrices, removing clusters with <5 members.

Using the above procedure, we obtained the following optimum number of clusters for each data set: 1. PR-IPL: 15 clusters (2 clusters with <10 members removed); 2. SZ-IPL: 12 clusters (1 cluster with <10 members removed); 3. PR-GCL: 13 clusters (7 clusters with <5 members removed); 4. SZ-GCL: 9 clusters (1 cluster with <5 members removed). Unshared-diagonal covariance matrices gave the optimal solution in all cases.

## Author contributions

MZ, JB, TY and TB designed the study, with input from all authors; MZ, JB and TY generated novel transgenic lines; FKJ developed custom 2-photon scan options; MZ and JB performed functional recordings and data pre-processing: population recordings around the eye, single cell recordings, comparison flat vs. natural stats noise stimulation (MZ), population recordings in the SZ (JB); JB performed paGFP experiments and retinal cell counts and data pre-processing; TY computed cell-density projections into visual space; PAR performed clustering analyses; TB performed additional data analysis and wrote the manuscript, with input from all authors.

## Acknowledgements

We thank Daniel Osorio and Katrin Franke for critical feedback, Philipp Berens for advising on statistical analyses, and Maxime Zimmermann for providing the 3D zebrafish model. The authors would also like to acknowledge support from the FENS-Kavli Network of Excellence and the EMBO YIP.

