## Supplementary Materials for "What the Zebrafish’s Eye Tells the Zebrafish’s Brain: Retinal Ganglion Cells for Prey Capture and Colour Vision"

### Supplementary Discussion

#### Population imaging from RGC dendrites and somata in the live eye

In this study, we used 2-photon imaging of Islet2b:mGCaMP6f signals in the eye's GCL and IPL to functionally survey RGC functions in larval zebrafish. While this approach likely provides for a useful approximation of what the zebrafish's eye tells the zebrafish's brain, two main caveats must be considered:

First, while Islet2b is an effective and popular marker for zebrafish RGCs (Janiak et al., 2019; Thisse et al., 2004), it is neither exclusive to RGCs nor inclusive of all RGCs. In our Islet2b:mGCaMP6f line, immunostaining against GFP and GABA revealed that some dACs also express GCaMP6f (Supplementary Fig. S1B), indicating that our dataset contains a minority of signals from dACs. In addition, small numbers of INL somata are labelled, indicating that also a minority of ACs contribute to our dendritic signals (AC somata are not included since these are easily discarded based on location). Conversely, not all axonal arborisation fields (AFs) in the brain, as revealed after DiO injection into the eye, were also strongly innervated by mGCaMP6f expressing RGCs (Supplementary Fig. S1C,D), suggesting that a subset of RGC types may be absent in our dataset. Finally, also a small fraction of central neurons were labelled as evident from their soma locations near the (pre)tectal neuropils.

Second, population imaging of RGC dendrites in the eye is potentially fraught with many of the same problems that are associated with delineating their axonal signals in the brain (Nikolaou et al., 2012). Specifically, the high density and overlap of dendritic processes across the IPL means that it is impossible to tell if groups of dendritic ROIs belong to the same RGC (Figs. 2D,E). Functional dendritic data is therefore mainly indicative of the local computations that occur within RGC dendrites as they integrate signals from BCs and ACs in different layers of the IPL and in different positions of the eye.

In contrast, the somata of RGCs in the GCL could generally be well segmented. In view of their strategic position next to the axon hillock, data from RGC somata may thus serve as the main indication of the signal sent from the eye to the brain. Nevertheless, addressing how exactly somatic calcium signals are linked to spikes sent down the optic nerve will be important in the future. This may then also go partway to explaining the marked reduction in Off-responses in somatic data compared to dendrites (Fig. 3B-D, Fig. 5A, Fig. 7B-D), and more broadly to drive our understanding of how this tiny animal's eye communicates with its brain.

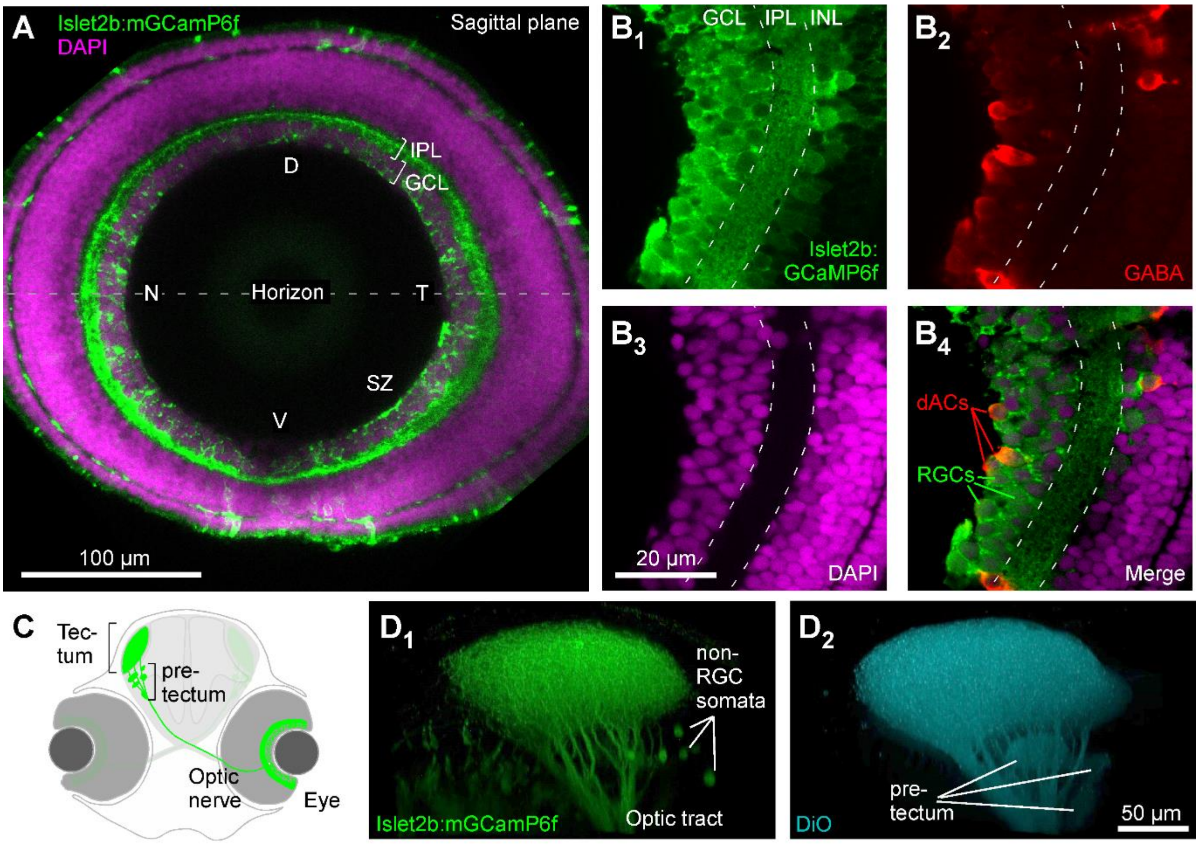


***Supplementary Figure S1 | related to Figure 2. Islet2b:mGCaMP6f expression in the larval zebrafish eye and brain. A,*** *7 dpf larva whole-eye sagittal plane confocal image of Islet2b:GCaMP6f expression (green) on the background of a DAPI stain, labelling all somata (magenta). D, dorsal; T, temporal; V, ventral; N, Nasal; SZ, Strike zone; INL Inner nuclear layer; GCL, Ganglion cell layer.* ***$B_{1-4}$,*** *Example higher magnification as in (A) from a second animal, with additional immunolabelling for GABA (red) to reveal GABAergic dACs and AC. Note the subset of somata showing both GABA labelling and mGCaMP6f expression ($B_4$). Note also the near doubling of GCL thickness across the region from the ventral retina (bottom) leading into the SZ (top).* ***C****, schematic of one eye's RGC projections in larval zebrafish into the brain (green).* ***D,*** *confocal projections of mGCaMP6f signal in the brain ($D_1$) and counter labelling by DiO injection in the eye ($D_2$). Though generally similar, $D_1$ shows expression in small numbers of brain-somata, while $D_2$ shows stronger labelling in pre-tectal axonal arborisation fields.*

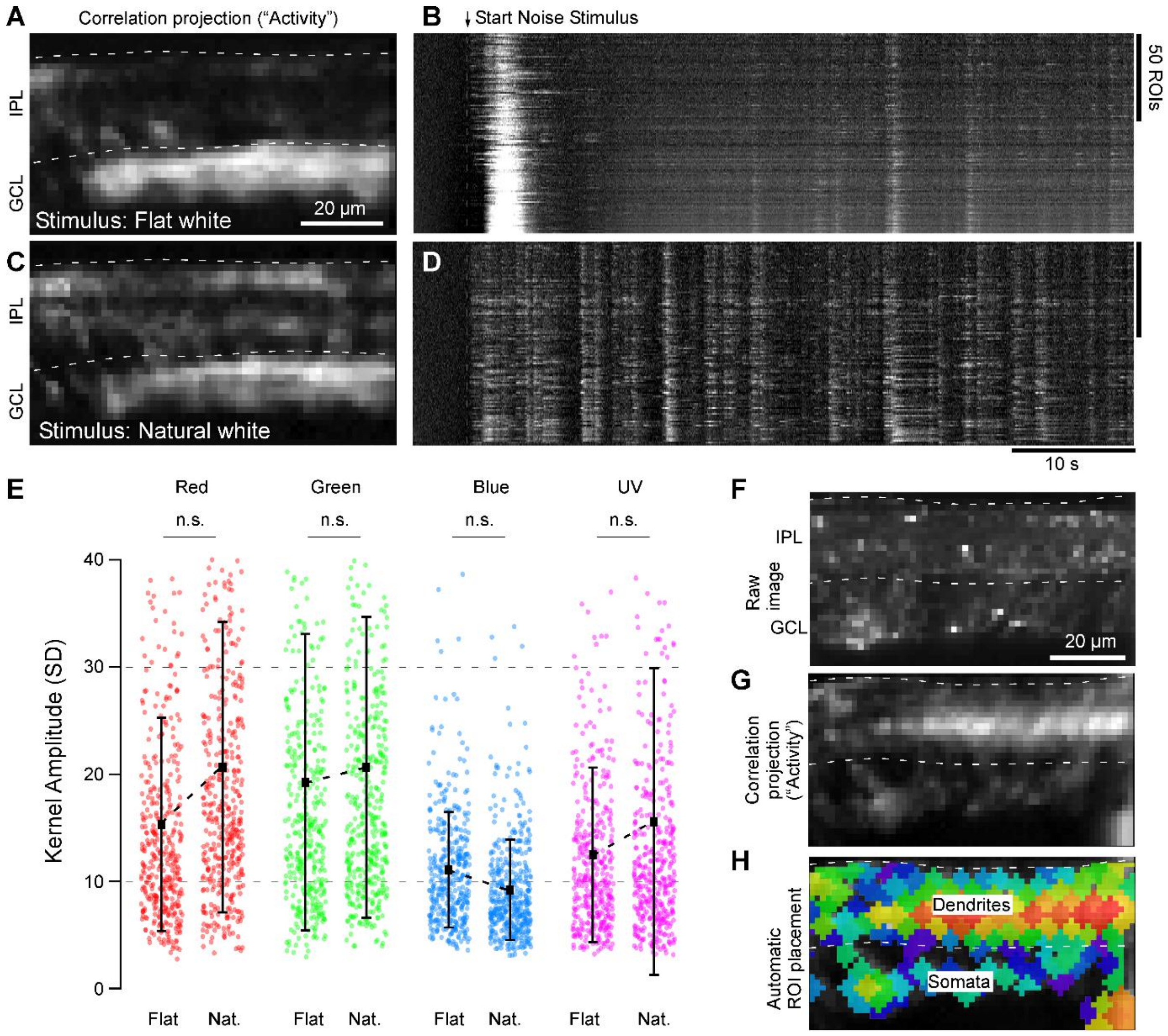


***Supplementary Figure S2 | related to Figure 2. Stimulation with 'natural white' light and ROI placement. A-D,*** *comparison of light-evoked activity in the same scan region in the SZ during stimulation with spectrally-flat white-noise (A, B) and identical sequence 'natural spectrum' white noise (i.e. with G, B and UV attenuated relative to R, cf. Fig. 2B) (C, D). Panels A,C show the correlation projection for the entire scan, while B, D shows a heatmap of all extracted ROIs from each scan, respectively. Note the strong initial response upon switching on the noise sequence in the flat-white condition (B), followed by a profound period of response suppression. In contrast, ROIs during the natural-white condition responded briskly to the noise sequence without showing strong adaptation (D). Similarly, a more diverse set of scan-regions strongly responded in the natural white condition (C) compared to flat-white (A). E, The mean of the distributions of resultant kernel amplitudes across n=6 such scans from an identical number of animals (n=388 and 428 ROIs for the flat and natural-white condition, respectively) were indistinguishable (Wilcoxon Rank Sum Test, 2 tailed). Based on these results, we decided to use natural-white noise stimulation throughout this study. F-H, example scan demonstrating typical automated ROI placement. The scan was manually segmented into IPL (F, top) and GCL (bottom). In parallel, we computed mean correlation over time between all pairs of neighbouring pixels for the entire scan (G), and the resultant correlation-projection image was in turn used to seed and flood-fill ROIs. For further details and a discussion about the rationale of this approach, see (Franke et al., 2017).*

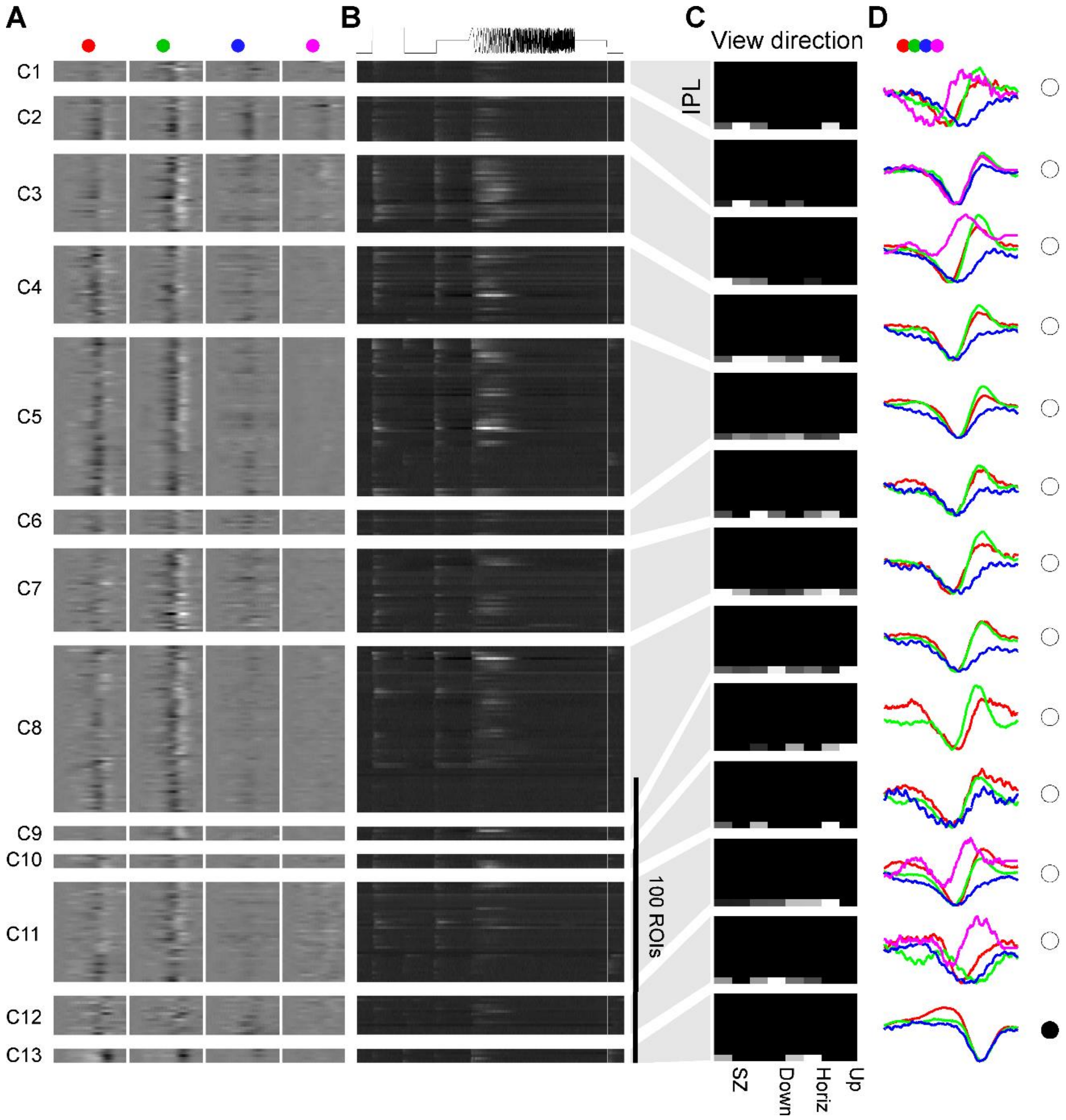


***Supplementary Figure S3 | related to Figure 6. Clustering of RGC somatic functions across the eye. A-D**, Somatic data from across the eye clustered based on spectral kernels, presented following the same organisation as used for dendritic data (Fig. 6A-C, E,F).*

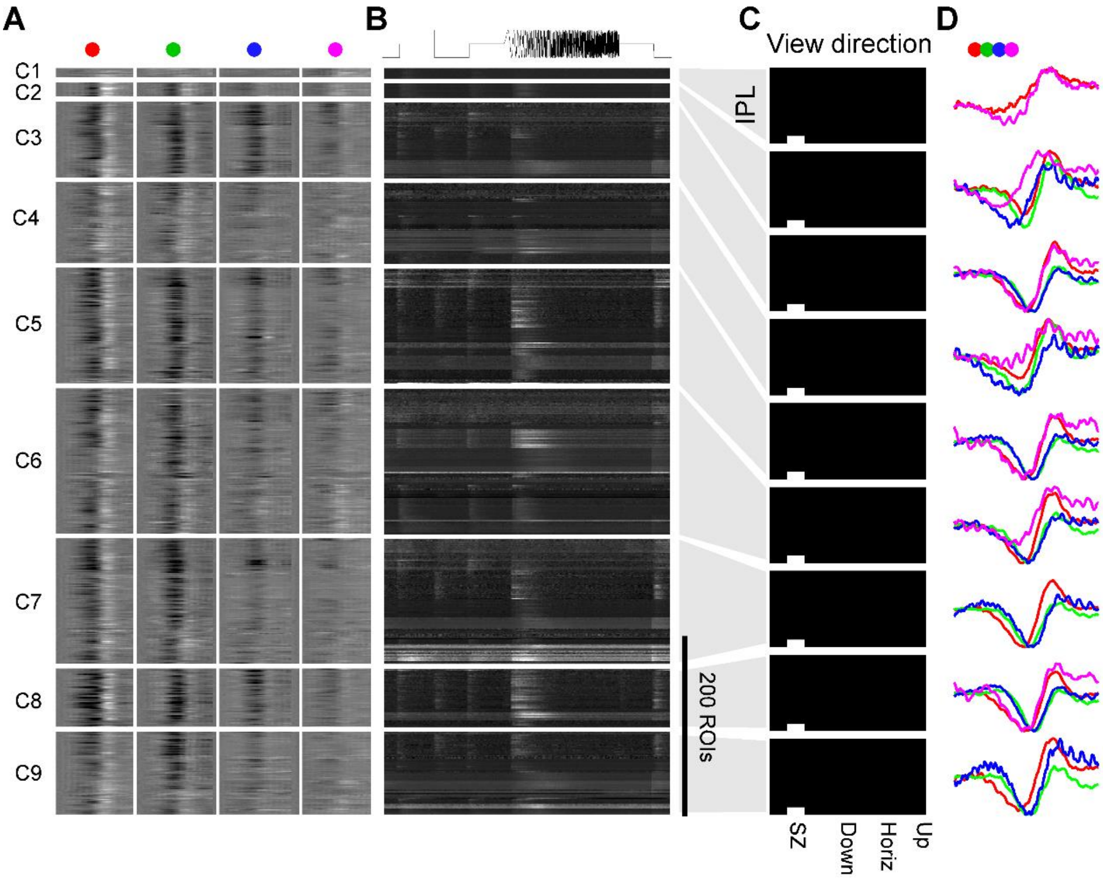


***Supplementary Figure S4 | related to Figure 8. Clustering of RGC somatic functions from the SZ***. ***A-D***, *Somatic data from the SZ based on spectral kernels, presented following the same organisation as used for dendritic data (Fig. 8A-C, E,F).*

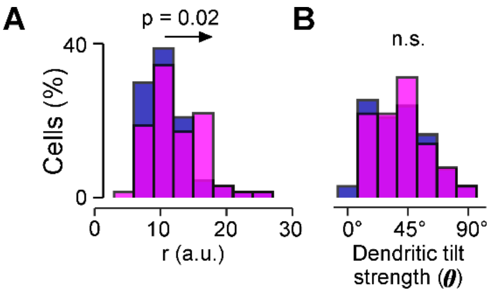


***Supplementary Figure S5 | related to Figure 9. Dendritic tilt, additional parameters***. ***A,B***, *Summary histograms of the distributions of r and θ (cf. Fig. 9D) for quantifying dendritic tilt in photo-labelled RGCs, pink: SZ, blue: Nasal. The distribution of r was weakly but significantly right-shifted in SZ RGCs relative to nasal RGCs (A), while the corresponding distributions of φ were non-statistically distinct (B). Both: Two-sample Kolmogorov-Smirnov test.*

### Supplementary Video Legends

***Supplementary Video S1 | related to Figure 2.*** *Background-subtracted but otherwise "raw" fluorescence responses of the example recording summarised in Figure 2. RGC dendrites (top half) and somata (bottom half) respond to the presentation of full-field tetrachromatic noise stimulation. Video plays at real time.*

***Supplementary Video S2 | related to Figure 4A-E.*** *Average Eye(x)-IPL(y) response over time across our entire dataset to an achromatic step of light, as shown in Fig. 4D ($t_{1,2}$) and E (left panels), starting with the Off-response (responses in the top of the IPL), followed by the On-response (bottom of the IPL). "Hot" colours indicate increased activity. Video plays in real time.*

***Supplementary Video S3 | related to Figure 4A-E.*** *As Supplementary Video S2, but for the temporal flicker portion of the stimulus ($t_{3,4}$). "Hot" colours indicate increased activity. Video plays in real time.*

***Supplementary Video S4 | related to Figure 4F-L.*** *As Supplementary Video S2, but instead of showing the mean step/flicker responses to achromatic stimulation, showing the average temporal kernels recovered from tetrachromatic stimulation (cf. Fig. 4I). From top left to bottom right: Red, Green, Blue, UV. Stronger colours indicate deviations above baseline. For clarity, approximately equal and opposite deviations below baseline are masked in this colour map and appear black. Video plays at 25% real time.*
